# Bulk and single-cell gene expression profiling of SARS-CoV-2 infected human cell lines identifies molecular targets for therapeutic intervention

**DOI:** 10.1101/2020.05.05.079194

**Authors:** Wyler Emanuel, Mösbauer Kirstin, Franke Vedran, Diag Asija, Gottula Lina Theresa, Arsie Roberto, Klironomos Filippos, Koppstein David, Ayoub Salah, Buccitelli Christopher, Richter Anja, Legnini Ivano, Ivanov Andranik, Mari Tommaso, Del Giudice Simone, Papies Jan Patrick, Müller Marcel Alexander, Niemeyer Daniela, Selbach Matthias, Akalin Altuna, Rajewsky Nikolaus, Drosten Christian, Landthaler Markus

**Affiliations:** Berlin Institute for Medical Systems Biology, Max-Delbrück-Center for Molecular Medicine in the Helmholtz Association, Hannoversche Str 28, 10115 Berlin, Germany; Institute of Virology, Charité-Universitätsmedizin Berlin and Berlin Institute of Health, Charitéplatz 1, 10117 Berlin, Germany; Max-Delbrück-Center for Molecular Medicine in the Helmholtz Association, Robert-Rössle-Strasse 10, 1312, Berlin, Germany; Department of Pediatrics, Charité – University Hospital Berlin, 13353 Berlin, Germany; Core Unit Bioinformatics, Berlin Institute of Health, Charité – University Hospital Berlin, 10117 Berlin, Germany; IRI Life Sciences, Institut für Biologie, Humboldt Universität zu Berlin, Philippstraße 13, 10115, Berlin, Germany

## Abstract

The coronavirus disease 2019 (COVID-19) pandemic, caused by the novel severe acute respiratory syndrome coronavirus 2 (SARS-CoV-2), is an ongoing global health threat with more than two million infected people since its emergence in late 2019. Detailed knowledge of the molecular biology of the infection is indispensable for understanding of the viral replication, host responses, and disease progression. We provide gene expression profiles of SARS-CoV and SARS-CoV-2 infections in three human cell lines (H1299, Caco-2 and Calu-3 cells), using bulk and single-cell transcriptomics. Small RNA profiling showed strong expression of the immunity and inflammation-associated microRNA miRNA-155 upon infection with both viruses. SARS-CoV-2 elicited approximately two-fold higher stimulation of the interferon response compared to SARS-CoV in the permissive human epithelial cell line Calu-3, and induction of cytokines such as CXCL10 or IL6. Single cell RNA sequencing data showed that canonical interferon stimulated genes such as IFIT2 or OAS2 were broadly induced, whereas interferon beta (IFNB1) and lambda (IFNL1-4) were expressed only in a subset of infected cells. In addition, temporal resolution of transcriptional responses suggested interferon regulatory factors (IRFs) activities precede that of nuclear factor-κB (NF-κB). Lastly, we identified heat shock protein 90 (HSP90) as a protein relevant for the infection. Inhibition of the HSP90 charperone activity by Tanespimycin/17-N-allylamino-17-demethoxygeldanamycin (17-AAG) resulted in a reduction of viral replication, and of TNF and IL1B mRNA levels. In summary, our study established in vitro cell culture models to study SARS-CoV-2 infection and identified HSP90 protein as potential drug target for therapeutic intervention of SARS-CoV-2 infection.

## Introduction

Diseases caused by coronaviruses (CoVs) range from asymptomatic and mild infections of the upper respiratory tract to severe acute respiratory distress, when the lower respiratory tract is infected. In addition to the six previously-known CoVs affecting humans, a novel CoV termed severe acute respiratory syndrome coronavirus 2 (SARS-CoV-2) has recently emerged. The novel SARS-CoV-2, which causes coronavirus disease 2019 (COVID-19), is still an ongoing global health threat since the beginning of the outbreak in late 2019 and has, at the time of writing this text, infected more than three million people worldwide [1]. The SARS-CoV-2 life cycle initiates with the attachment of the virion to the cell surface and subsequent binding to the angiotensin converting enzyme 2 (ACE2), followed by proteolytic cleavage and internalization [2-4]. Non-structural proteins are then translated to form a replicase-transcriptase complex (RTC), in which the full genomic RNA, as well as subgenomic RNAs are generated within double membrane vesicles (DMV) [3-5]. Incoming viral RNA is detected by sensors such as IFIH1 (interferon induced with helicase C domain 1; also known as MDA5) and DDX58 (DExD/H-Box helicase 58; also known as RIG-I), which trigger the antiviral response. This sensing and signaling is impaired by a range of viral factors, e.g. replication within DMVs, RNA capping and methylation, or shortening of the poly-U tail on the minus strand RNA [6, 7]. Furthermore, inhibition of IRF activity [8] and a delayed induction of interferon-stimulated genes (ISGs) compared to Influenza virus infection or type I interferon treatment itself [9] was observed in SARS-CoV infection. Importantly, accessory genes in the SARS-CoV genome, like ORF6, may code for antagonists of interferon signaling [10].

Following production of subgenomic RNAs, during which a constant 5’ leader is prepended by a process called discontinuous transcription [11], the viral genes are translated either in the cytoplasm (nucleocapsid protein, N), or at the endoplasmic reticulum (ER; envelope (E), membrane (M), spike (S) and open reading frame 3 (ORF3) proteins) [12, 13]. The substantial increase in ER translation causes ER stress, which triggers the unfolded protein response. This is then in turn integrated with double stranded RNA sensing at the level of eukaryotic initiation factor 2 alpha (eIF2alpha) phosphorylation [14]. The ER stress response is likely attenuated by the viral E protein [15, 16]. Accordingly, heat shock proteins (HSPs), which ameliorate ER stress, have been described to be generally relevant in virus infections [17]. Furthermore, ER stress induces autophagy, a cell recycling pathway which can be used by some viruses for productive replication [18]. Finally, dysregulation of microRNA (miRNA) expression and subsequent alterations in gene expression patterns have also been reported to play a role in infected cells [19, 20].

Comprehensive profilings of SARS-CoV-2-mediated gene expression perturbations are just beginning. A recent in-depth analysis of the transcriptional response to SARS-CoV-2 in comparison to other respiratory viruses in cells and animal models revealed a virus-specific inflammatory response [21]. Of particular interest are methods such as single-cell RNA-sequencing (scRNA-seq), which allow the characterization of heterogeneity over the course of infection, which may be masked at the population level [22-27], but also small RNA sequencing, which reveals miRNAs and other small RNAs [28, 29].

Here, we performed a comprehensive analysis of three human cell lines infected with SARS-CoV or SARS-CoV-2, namely the gut cell line Caco-2, the lung cell lines Calu-3, and H1299. We generated scRNA-seq, poly(A)^+^ and total RNA transcriptomic data as well as small RNA profiling in infection time courses for both viruses.

Efficiency and productivity of infection as well as the interferon response was remarkably different between the cell lines. Interestingly, SARS-CoV-2 induced a two-fold higher expression of interferon stimulated genes (ISGs) than SARS-CoV. In addition, we found strong induction of miR-155 with both viruses, suggesting a role for this miRNA in the progression of infection. The scRNA-seq data showed that, while canonical ISGs such as interferon induced protein with tetratricopeptide repeats 1 and 2 (IFIT1/IFIT2) were broadly induced, interferon beta (IFNB1) was expressed only in a subset of infected cells. Furthermore, the transcriptional induction of nuclear factor-κB (NF-κB) targets could be temporally separated from the interferon gene induction. Detailed investigations of cellular gene expression programs suggest an involvement of the protein folding chaperone and autophagy regulator HSP90 in the viral infection cycle. Inhibition of HSP90 by 17-N-allylamino-17-demethoxygeldanamycin (17-AAG) resulted in reduced viral replication and TNF and IL1B mRNA levels. Overall, our study provides a detailed picture of the gene expression changes in cell line models for CoVs and particularly SARS-CoV-2, highlights the cell-type specificity of the transcriptional response to infection, and identifies potential targets for therapeutic interventions.

## Results

### Different permissiveness of SARS-CoV-2 infection in cell lines

To establish cell culture systems for studying SARS-CoV-2 replication and host cell responses, we examined the epithelial lung cancer cell lines, H1299 and Calu-3, as well as the epithelial colorectal adenocarcinoma cell line, Caco-2, which is frequently used as a coronavirus cell culture model [30-32]. Transfection of poly-I:C RNA, resulted in induction of IFIT1, IFIT2 and OAS2 (2’-5’-Oligoadenylate Synthetase 2) genes in Calu-3 and H1299 cells, indicating sensing of cytoplasmic foreign RNA is active in these cell types. This response was not observed in Caco-2 cells (Fig. S1A), which only poorly expresses viral RNA receptor genes, IFIH1/MDA5 and DDX58/RIG-I (Supplementary data 1).

For all cell lines, we performed a comprehensive analysis of transcriptome changes at different time points post infection with SARS-CoV (Frankfurt strain) and SARS-CoV-2 (patient isolate BetaCoV/Munich/BavPat1/2020|EPI_ISL_406862) at an MOI of 0.3 (Fig. 1A, Supplementary Table 1). The percentages of viral transcripts in intracellular RNAs as determined by Poly(A)^+^ and RNA-seq sequencing were low in H1299 cells for both viruses in contrast to Caco-2 and Calu-3 cells (Fig. 1C, Supplementary Table 2). Accordingly, the yield of infectious virus particles was higher for the permissive cell lines Caco-2 and Calu-3 (Fig. S1A). The low susceptibility of H1299 cells might be attributed, at least partially, to the low expression of the SARS-CoV receptor ACE2, based on RNA-sequencing data and Western blot analysis (Fig. 1B).

**Fig. 1.**
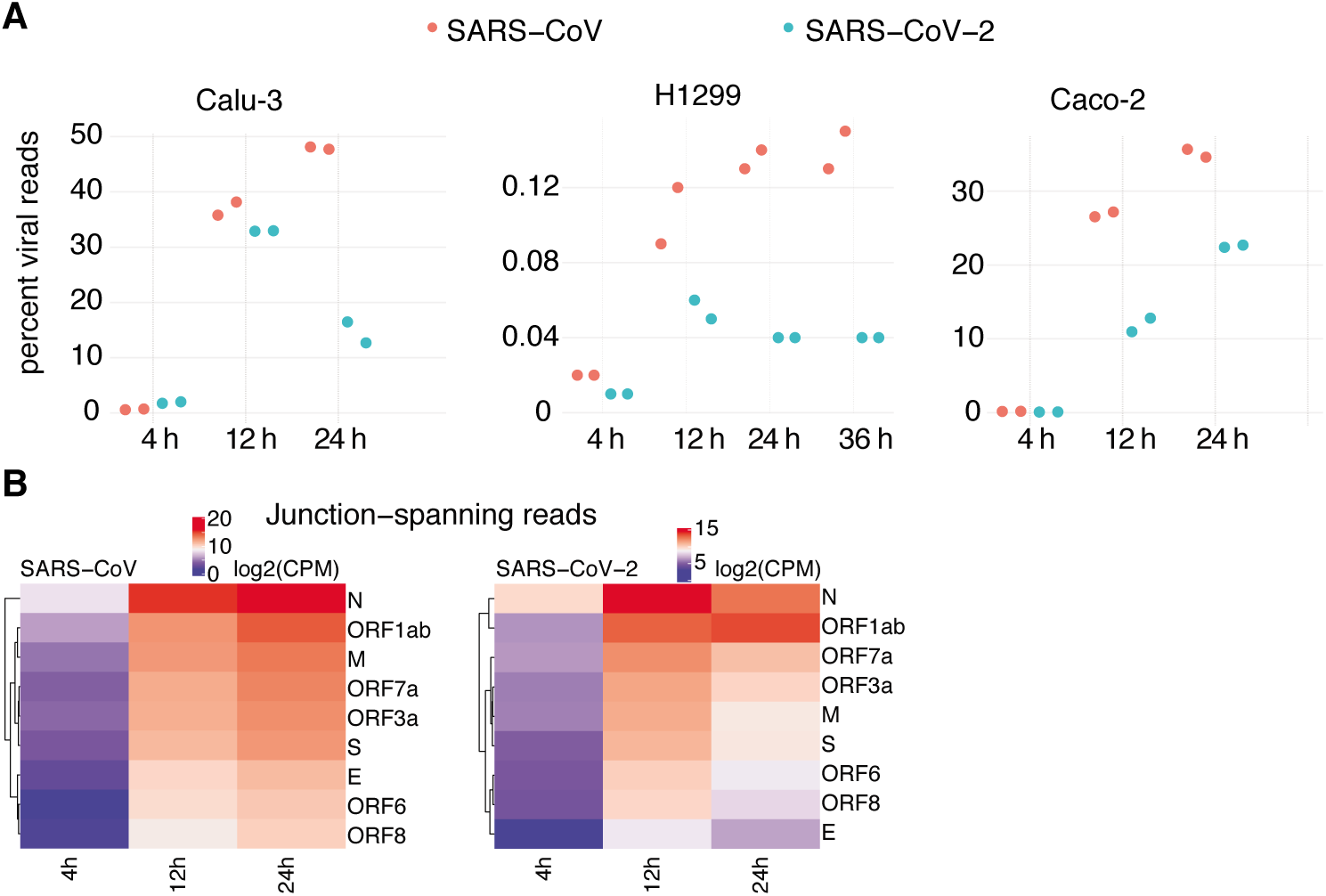
**A**, Viral read percentages of total reads of the respective cell lines at different time points of infection. All cell lines show an induction as a function of time. H1299 show the lowest percent-age of viral reads. SARS-COV infected cells exhibit the highest amount of viral reads. **B**, Heatmaps of canonical junction-spanning viral reads, averaged across biological replicates per time point, expressed in TMM-normalized counts per million.

By counting poly(A)^+^ or total RNA-seq reads spanning the junction of the viral leader and its downstream gene (82.6% of virus-mapping split reads), we accurately quantified the relative amounts of subgenomic viral mRNAs [33, 34]. We observed a consistent hierarchy of gene expression across time, mostly dominated by viral mRNAs encoding the M gene (Fig. 1D, Fig. S1D), similar to a recent report for the alpha Human CoV-229E (HCoV-229E) [35]. At later time points post infection, the relative amount of ORF7a generally increased (Figure S1E). Notably, this approach failed to detect expression of leaders immediately adjoining ORF7b or ORF10 (Supplementary Table 3).

By visual inspection, Caco-2 cells appear hardly affected by the infection; whereas, Calu-3 clearly show signs of impaired growth and cell death at 24 hours post infection (hpi), particularly when infected with SARS-CoV-2 (Fig. S1F). Taken together, we show that the three infected cell lines show distinct responses in respect to the course of SARS-CoV/-2 infection.

### SARS-CoV-2 infected Calu-3 cells show a strong induction of interferon-stimulated genes

Although the infection is comparable in Caco-2 and Calu-3 cells, judging based on the amount of intracellular viral RNA and virion yield (see above), the host transcriptome responses were markedly different (Fig. 2A, 2B). In case of the SARS-CoV-2-infected Caco-2 cells, an increase in expression of a number of genes (activating transcription factor 3, ATF3; early growth response 1, EGR1; immediate early response 3, IER3;) was detected that are typically activated in response to ER stress. In contrast, Calu-3 cells showed a strong increase in expression of ISGs such as the IFIT1, IFIT2, and interferon genes. This response was absent in Caco-2 cells likely due to the reduced expression of pattern-recognition receptors, IFIH1 and DDX58 that activate transcription of target genes via interferon regulatory factors (IRF) and NF-κB signaling [36, 37], which supports similar results for SARS-CoV infection [38] (Supplementary data 1).

**Fig. 2.**
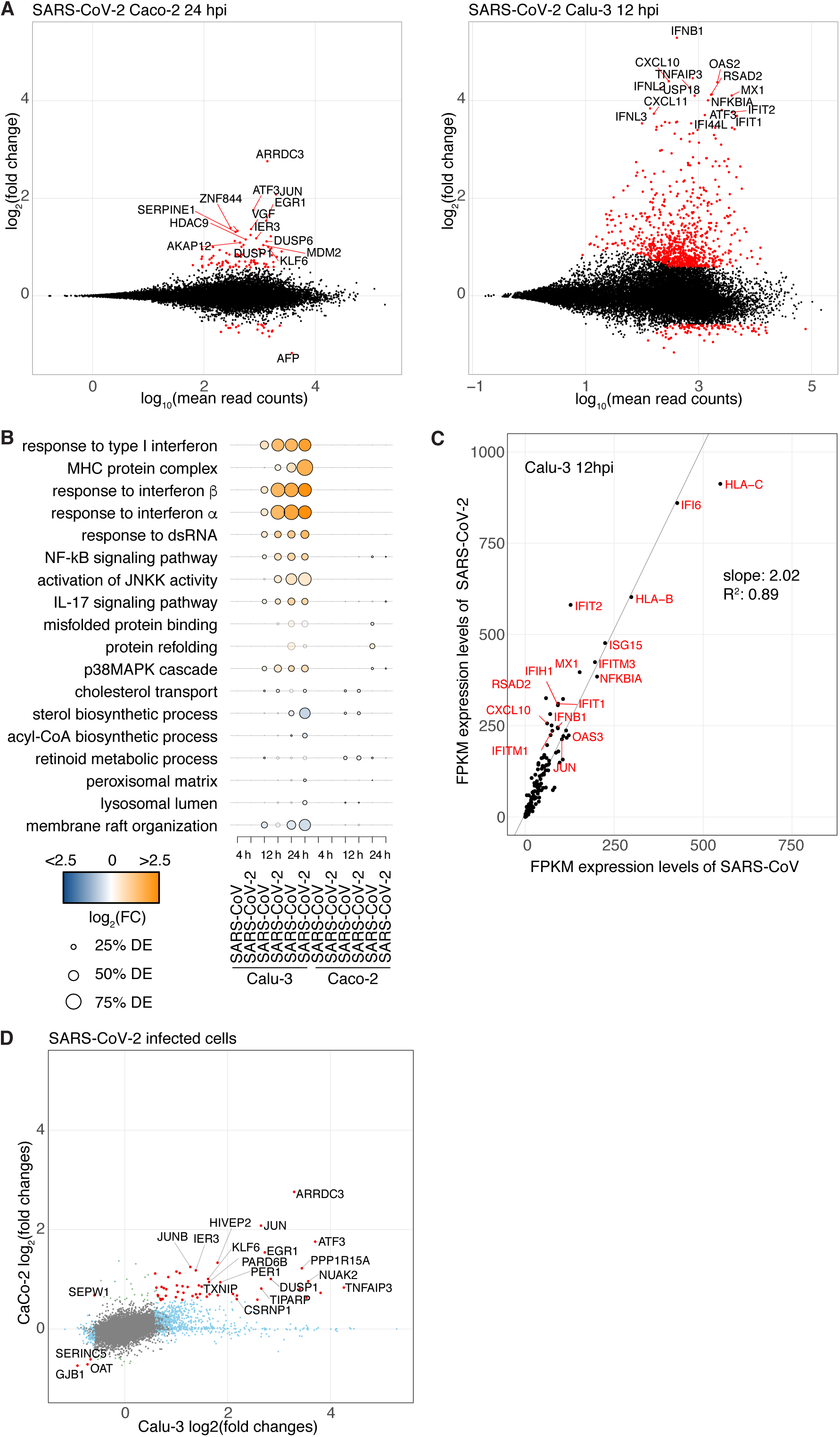
**A**, MA plots of differentially expressed (DE) genes at the mRNA level of Calu-3 (12 hpi) and Caco-2 cells infected with SARS-CoV-2. X-axis depicts log10 of mean counts associated with a gene. Y-axis depicts mRNA log2 fold changes. Significant differentially expressed genes (red dots) are defined as those with an absolute log2 fold change of greater than 0.58 and an adjusted p-value > 0.05. Selected outliers are labeled. **B**, Gene Ontology (GO) term enrichment for different comparisons. A sample of gene sets overrepresented significantly in at least one comparison (hypergeometric test, adjusted p-value < 0.05). Dot size depicts percentage of gene set members classed as differentially expressed in one comparison. Gene sets overrepresented significantly are displayed as solid dots. Dot colour represents the average log2 fold change of mRNAs of a gene set in that comparison. Calu-3 cells exhibit a higher DE than CaCo-2 cells challenged by infection. **C**, expression values (FPKM) of genes significantly induced in cells infected with either viruses at 12 hpi in Calu-3 cells. **D**, Genes exhibiting significant changes both cell lines are shown in red, log2-transformed fold changes of SARS-CoV-2 infected cells at 12 hpi (Calu-3, horizontal axis) and 24 hpi (Caco-2, vertical axis). Genes exhibitingsignificant changes in both cell lines are shown in red, significant changes only in Calu-3 cells in light blue, only in Caco-2 cells in light green. All other genes are shown in grey. Selected genes with significant fold changes are labeled.

When comparing the expression of ISGs in virus-infected Calu-3 cells, our data revealed that the expression levels of ISGs were on average about twice as high for SARS-CoV-2 compared to SARS-CoVinfected cells (Fig. 2C) at similar amounts of viral RNAs present in the cells (Fig. 1C, right panel). This difference in the extent of the ISG response may be of clinical relevance, since cytokines [39] are among the induced ISGs (Fig. S2B), and their expression might be connected with pathologies such as the acute respiratory distress syndrome (ARDS) in CoV infections [40, 41].

Next, we compared the induction of ISG in SARS-CoV-2 between experiments, and with a recently published gene expression study of normal human bronchial epithelial (NHBE) cells, A549 cells with and without ACE2 expression, and Calu-3 cells upon infection with SARS-CoV-2 [21] (Supplementary Table 2, Fig. S2B). Whereas infection of Calu-3 cells showed strong reproducibility across experiments from our study (Fig. S2C) and across labs (Fig. S2D), there were remarkable differences between the results of Calu-3 on one side and A549 cells and NHBE cells on the other side, which both lack an induction of interferon beta/lambda genes (Fig. S2EF).

### Expression of ARRDC3 and TXNIP genes is induced independently of RNA sensing

To identify genes that might be altered independently of RNA sensor-triggered signal cascades, we compared gene expression changes between the Caco-2 and Calu-3 cell lines at 12 hpi with SARS-CoV-2 (Fig. 2D, Fig S2A). Two genes, arrestin-related domain-containing protein-3 (ARRDC3) and thioredoxin-interacting protein (TXNIP), stood out among the few that were significantly dysregulated upon infection with either viruses. Both genes encode proteins that are involved in regulation of signaling pathways [42]. ARRDC3 mediates G protein–coupled receptor lysosomal sorting and apoptosis-linked gene 2-interacting protein X (ALIX) ubiquitination [43]. ALIX is a Lys63-specific polyubiquitin binding protein that functions in retrovirus budding and Dengue virus propagation [44, 45]. TXNIP is involved in the regulation of glucose and lipid metabolism [46], and has been shown to be involved in initiation and perpetuation of NLRP3 (nucleotide-binding domain and leucine-rich repeat and pyrin domain containing 3) inflammasome activation [47, 48].

To conclude, most gene expression changes in response to SARS-CoV-2 infection are likely triggered by RNA sensors. However, there are a few exceptions, and the mechanism of induction, and the function of these proteins during virus replication remains to be elucidated.

### MicroRNA miR-155-3p is expressed in SARS-CoV and SARS-CoV-2 infected cells

In addition to assessing mRNA changes, we have also profiled small RNAs in the context of Calu-3 infections. Both viruses trigger a close to 16-fold upregulation of miR-155-3p, the “star” form, and an almost 3-fold upregulation of miR-155-5p (Fig. 3A, 3B, S3A). Importantly, the primary miRNA precursor gene, miR-155 host gene (MIR155HG), was also upregulated in polyA-seq and total RNA-seq data by about 10-fold, suggesting that the increase of two miRNAs was primarily driven by transcription (Fig. S3B).

**Fig. 3.**
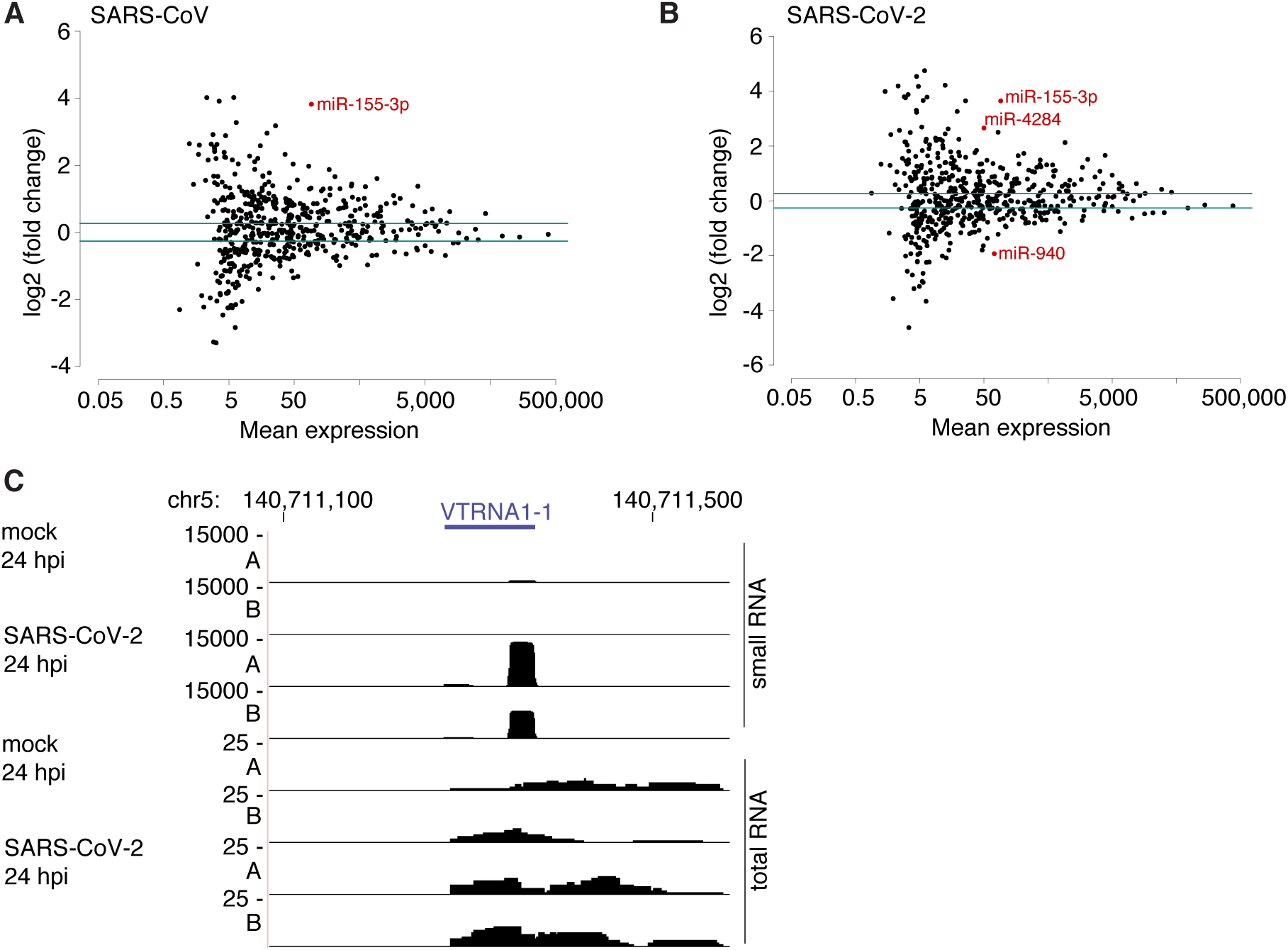
**A, B**, Scatterplots of miRNA mean expression versus log2-fold-changes testing for significant differences compared to mock long after the infection with SARS-CoV (A), and SARS-CoV-2 (B). miRNAs found differentially expressed are indicated in red. **C**, Coverage plot of the vaultRNA gene VRNAT1-1 of replicate A and B at indicted times after infection with or without SARS-CoV-2.

Interestingly, the miRNA profiling identified small RNAs mapping to vault RNA (VTRNA) genes (Fig. 3C, S3C). The function of vtRNAs has not been fully elucidated, but the mature VTRNA1-1 was recently discovered as a negative regulator of autophagy [49]. VTRNA-derived sRNAs can be processed by DICER and bound by Argonaute proteins [50, 51].

### Single-cell RNA-sequencing of SARS-CoV- and SARS-CoV-2-infected Calu-3 cells shows differences in interferon-stimulated gene expression

To assess gene expression changes on the level of individual infected Calu-3 cells, we performed scRNA-seq at different time points post infection for both SARS viruses. At 4 hpi, the number of cells bearing viral RNA was between 40% and 60% (Fig. S4A, supplementary table S4). At 8 hpi and 12 hpi, all cells contained viral RNA. The distribution of viral load (percentage of viral RNA per cell) was comparable for the two viruses, and showed the expected increase from 4 hpi to the later time points after infection (Fig. S4B).

The analysis of scRNA-seq data showed that cellular transcriptomes grouped by infection and type of virus (Fig. 4A, S4C). Two small groups made up of cells (cluster 13), derived from all time points and primarily, but not only SARS-CoV-2 infected cells showed high expression of the IFNB1 (Fig. 4B, 4D, S4D, S4E, S4F). As described above, we observed RNA sensing-independent expression of ARRDC3 in cells infected with either virus. ARRDC3, as well as the pro-apoptotic gene protein phosphatase 1 regulatory subunit 15A (PPP1R15A, also known as growth arrest and DNA-damage-inducible 34; GADD34), which had previously been related to ER stress [52], were highly expressed in the interferon gene expression cluster, but also in cells with very high levels of SARS-CoV-2 RNA (cluster 11; Fig. 4B, S4G). The mechanism leading to interferon expression in cluster 13 remains to be investigated.

**Fig. 4.**
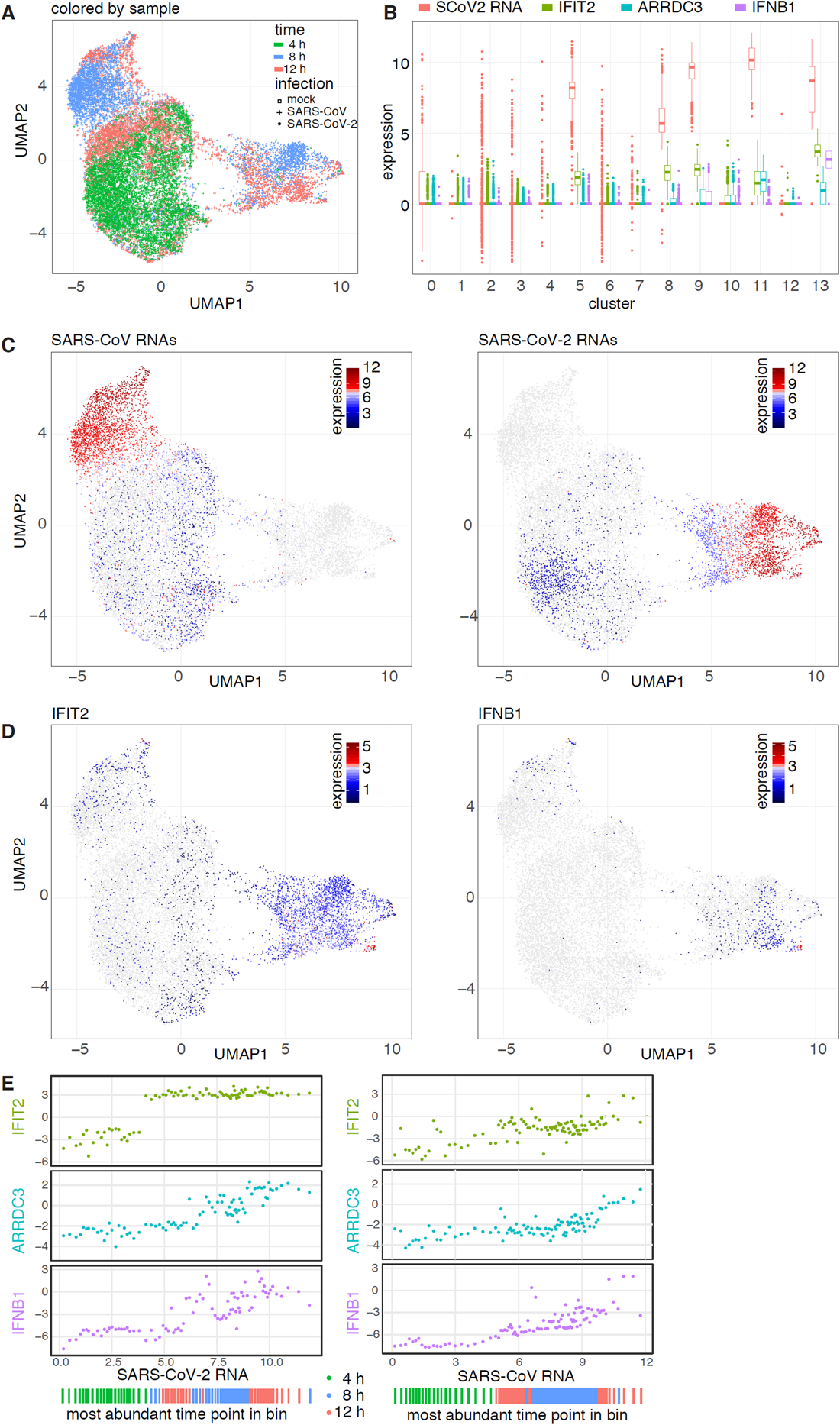
**A**, Based on scRNA-seq data cells are projected in two dimensions using Uniform Manifold Approximation and Projection (UMAP). **B**, Same projection, but cells are colored by SARS-CoV and SARS-CoV-2 viral transcript load. **C**, Distribution of expression values of selected viral and mRNA transcripts per cluster. **D**, Coloring by expression levels of IFIT2 (left) and IFNB1 mRNAs (right). **E**, Horizontal axis for each panel represents relative, log2 transformed levels of SARS-CoV-2 RNA (left) or SARS-CoV (right) per bin. The vertical axes in the panels represent relative, log2 transformed expression levels for the indicated gene per bin. Every dot represents a bin containing 50 cells. The most prominent time point per bin is indicated at the bottom.

In agreement with the bulk RNA-seq data, we observed a strong increase of expression of ISGs, IFIT1 and IFIT2, in infected cells, and particularly those exposed to SARS-CoV-2 (Fig. 4D, S4H). Likewise, MIR155HG, though poorly detected, resembled the expression pattern of IFIT1 and IFIT2 genes (compare Fig. S4I with 4B, and S4J with 4C/S4H).

To relate host gene expression to the accumulation of viral RNAs, cells were ordered by increasing amount of viral RNA, and arranged into bins of 50 cells. For each bin, the correlation with viral RNA was calculated for both viruses, indicating a strong relationship for the amount of viral RNA with ARRDC3 mRNAs (Fig. S5K). For selected genes, the expression level per bin was plotted against the amount of viral RNA (Fig. 5E). For IFIT2, this result indicated an likely off-on switch between 4 hpi and 8 hpi, with the expression levels being independent of the amount of viral RNA. For ARRDC3 however, the mRNA transcript numbers correlated well with the accumulation of viral RNA (R2 = 0.77). For the interferon beta gene IFNB1, a certain amount of viral RNA appeared to be a prerequisite for the IFNB1 mRNA levels, however beyond that threshold per-bin expression levels were rather variable. For SARS-CoV, the IFIT2 switch could not be observed, and the accumulation of ARRDC3 mRNA started only at higher levels of viral RNA (Fig. 4E).

**Fig. 5.**
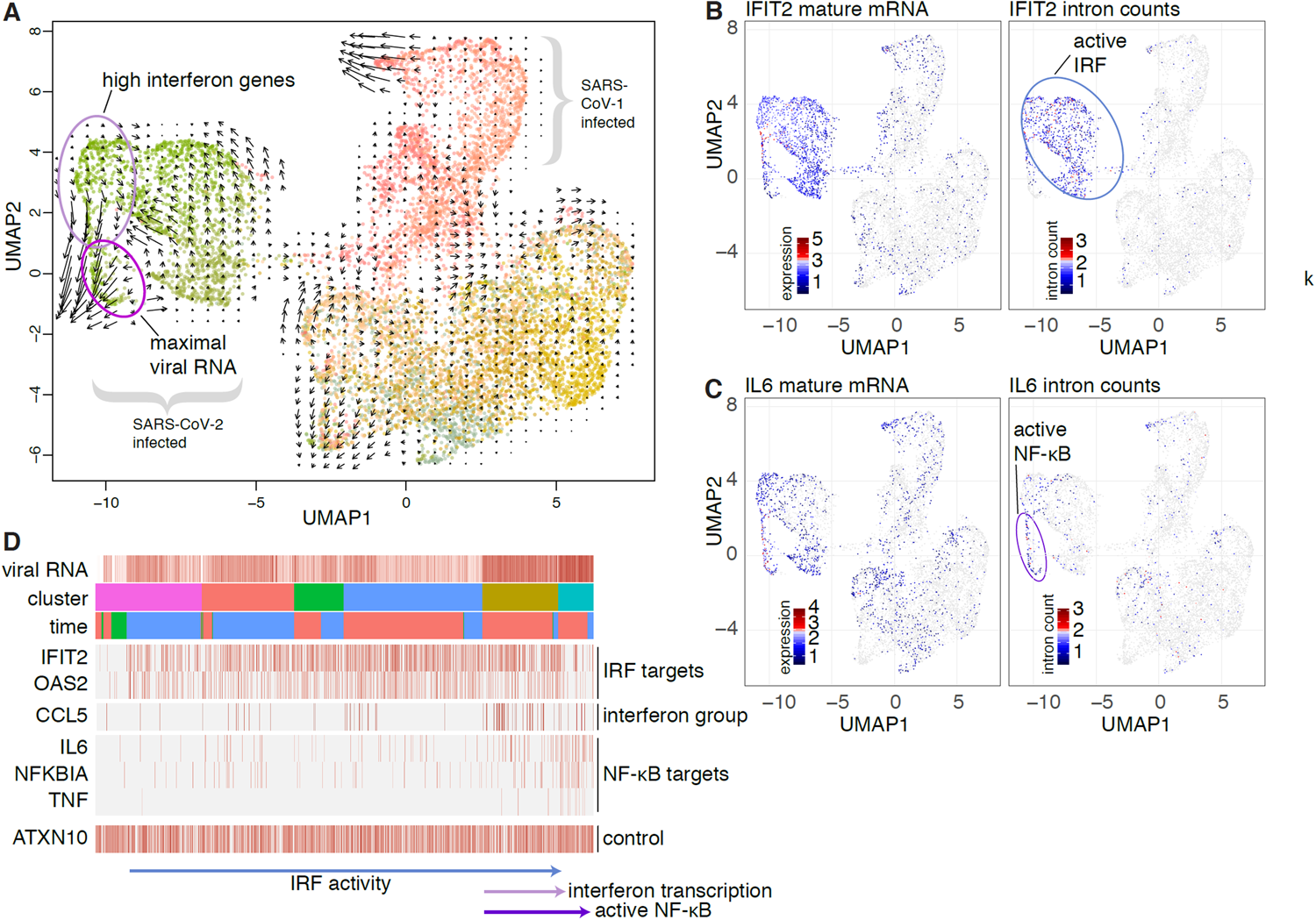
**A**, Cells were embedded into diffusion map space and by UMAP projected using 20 diffusion components into two-dimensional space. Areas of interest are marked. Arrows represent trajectories based on RNA velocity. **B, C**, Projection as in A, but colored by mature (left column) and intron-exon counts (right column) of IFIT2 (top) and IL6 (bottom). **D**, Columns in the heatmap represent cells. Plotted are, from top to bottom, the amount of SARS-CoV-2 viral RNA, the cluster ID, the harvesting time point, and intron counts for the indicated genes. Cells (columns) are first sorted by cluster, then by increasing amount of viral RNA.

In order to identify genes co-regulated with IFNB1, we performed a correlation analysis of cells binned by increasing IFNB1 and ARRDC3 expression level (Fig. S4L). Using this approach we found a putative co-regulation of the four interferon lambda genes (IFNL1-4), the chemokine genes, C-X-C Motif Chemokine Ligand 9 (CXCL9) and C-C Motif Chemokine Ligand 5 (CCL5), and the cholesterol-25-hydroxylase gene, CH25H. This enzyme, as well as its product 25-hydroxycholesterol, have been shown to act against a range of viruses [53]. In addition, two other genes were found in this group, the sodium voltage-gated channel alpha subunit 3 gene (SCN3A) and the dual oxidase 1 gene (DUOX1) (Fig. S4M), which are poorly expressed in cells outside cluster 13. Whereas SCN3A has previously not been described in the context of virus infections, DUOX1 promotes the innate immune defense to pathogens via the production of reactive oxygen species in mucosae [54].

For a better visualization of the effect of infection on cell clustering, we also analyzed the cells without taking viral transcripts into account, confirming stronger alterations of the cellular transcriptome in SARS-CoV-2 compared to SARS-CoV infections (Fig. S4N). Of note, the cells bearing interferon mRNAs clustered together independent of the virus.

### RNA velocity reveals transient induction of interferon genes and temporal resolution of NF-κB signaling

To better understand the nature of interferon gene induction in the context of CoV infection, we performed RNA velocity, which uses sequencing reads originating from introns to measure the amount of nascent mRNA [55]. This method applies additional filtering and embeddings, leading to different two-dimensional projections. This analysis showed a trajectory from cells expressing interferon and interferon-correlated genes to the cells with maximal amount of viral RNA but not expressing interferon genes (Fig. 5A, S5A, S5B). This finding suggests that induction of interferon genes is short and transient during viral replication. We observed that target genes of interferon regulatory factors (IRFs) such as IFIT2, IFIT1 or OAS2 [56] show high intron counts in an intermediate state during the accumulation of viral RNA (Fig. 5B, S5C), but this signal later decreases. At this time, however, NF-κB target genes such as interleukin 6 (IL6), tumor necrosis factor (TNF) or NF-κB inhibitor alpha (NFKBIA) [57] had still high intron counts (Fig. 5C, S5D). Taken together, this finding suggests that IRF-regulated genes are transcribed before NF-κB target genes, with IRF activity ceasing late (Fig. 5D). Interestingly, we also observed a minor increase in intronic counts for ACE2 transcripts in SARS-CoV-2 infected cells (Fig. S5E, S5F), suggesting a transcriptional activation of the viral receptor gene during infection, as observed recently [58].

### Single-cell RNA-sequencing of SARS-CoV and SARS-CoV-2 infected H1299 reveals a potential involvement of HSP90AA1 in the progression of infection

As shown above, transcriptional changes in bulk and scRNA-seq data from Calu-3 infected cells were dominated by the interferon response. In order to detect more subtle alterations in the cellular transcriptomes, we applied the scRNA-seq likewise to SARS-CoV/-2 infected H1299 cells, which are only partially permissive to the infection.

Despite the overall low amount of viral RNA in infected H1299 cells (Fig. 1C), the percentage of cells bearing viral transcripts was remarkably high (Fig. S6A, supplementary table S4), indicating that virions indeed are able to enter the cells. As seen in the bulk RNA-seq, transcriptional changes were subtle, and cells thus do not group into discrete clusters (Fig. S6B, S6C, S6D). When correlating individual genes with the amount of viral RNA over cells, we found a positive correlation with HSP90 alpha family class A member 1 (HSP90AA1) with the amount of SARS-CoV-2 RNA, but not SARS-CoV (Fig. 6A). Upon closer inspection, we observed higher expression of HSP90AA1 mRNA and RNA velocity values in cells with SARS-CoV-2 viral RNA compared to those without (Fig. 6B). We thus went back to the Calu-3 data and found a similar HSP90AA1 expression pattern in Calu-3 cells harvested at 4 hpi (Fig. 6C-E).

**Fig. 6.**
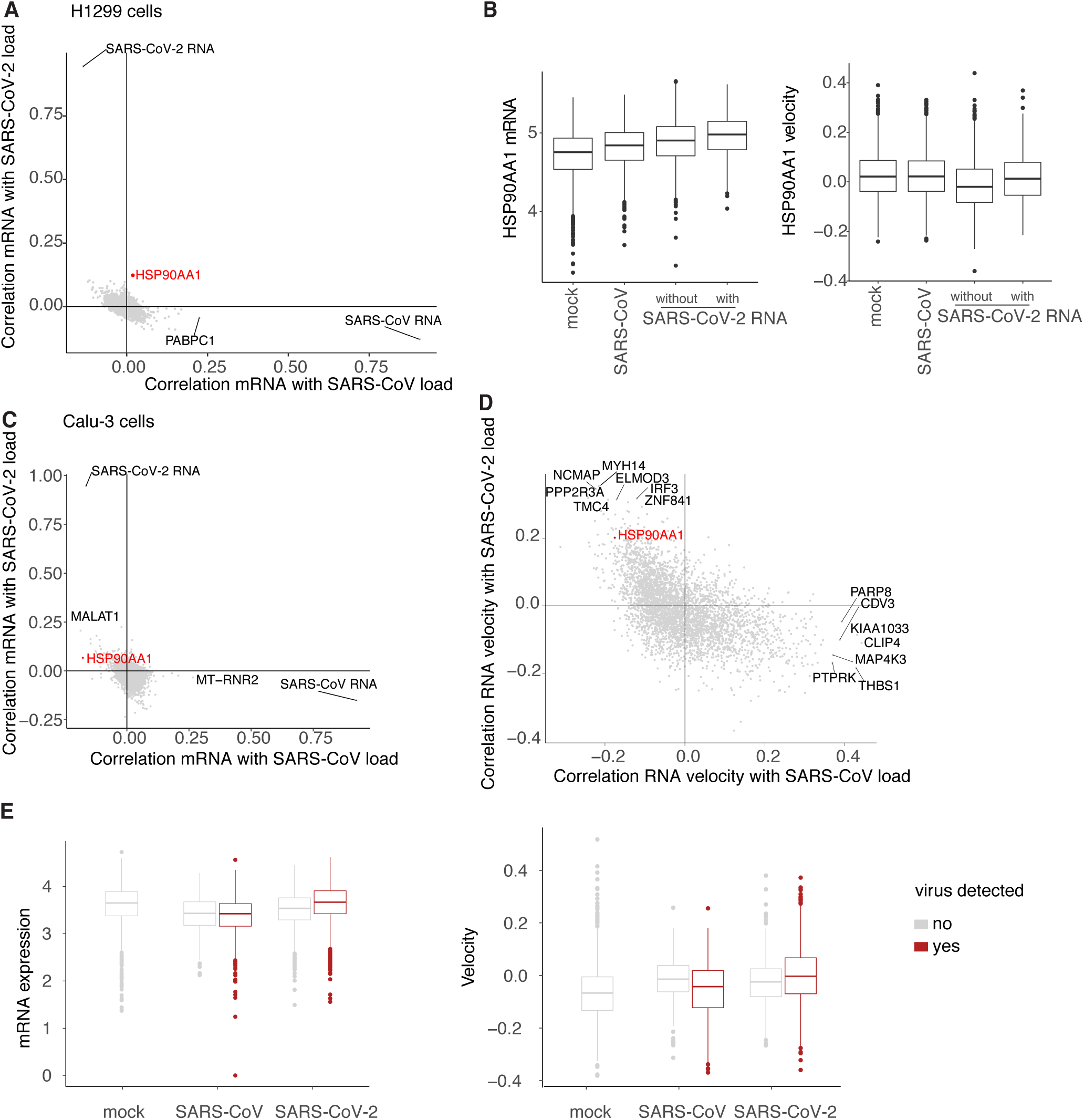
**A**, Correlation of gene expression values with the amount of SARS-CoV RNA (y-axis) and SARS-CoV-2 RNA (x-axis) in the H1299 scRNA-seq data. **B**, Distribution of HSP90AA1 mRNA (left) and RNA velocity (right) values in mock and SARS-CoV samples, and, for SARS-CoV-2 samples, split by cells with and without viral RNAs. **C**, As in A but for the Calu-3 4 hpi samples. **D**, As in C but with RNA velocity instead of mature mRNA values. **E**, Distribution of HSP90AA1 mRNA (left) and RNA velocity (right) values for mock, SARS-CoV and SARS-CoV-2 samples, split by cells with and without viral RNA.

### Inhibition of HSP90 reduces viral yield

The involvement of HSP90AA1, a highly-conserved molecular chaperone, in viral infections has since a long time been discussed to be involved in the infection of a range of viruses [59]. In order to explore the effect of HSP90 on SARS-CoV-2 replication in Calu-3 cells, we applied the HSP90 inhibitor 17-AAG to cells one hour after viral absorption and measured virus yield and RNA in the supernatant after 16 hpi (Fig. 7A) and 8 hpi (Fig. 7B). For the 8hpi experiment, also intracellular RNA was measured (Fig. 7B, bottom panel). At a concentration of 800 nM, 17-AAG reduced the viral yield to about 50%. In an additional experiment with similar outcomes (Fig. 7C, 7D), we assessed changes in host cell mRNA expression of infected cells to treatment with 17-AAG (Fig. 7E). Interestingly, whereas IFIT2 mRNA expression levels were only slightly reduced in cells treated with the inhibitor 17-AAG, reduction in mRNA levels for PPP1R15A and INFL1, and particularly for IL1B and TNF, were much more pronounced. This result could point to a complex regulation of these genes involving HSP90 activity. The treatment with 17-AAG had no apparent effect on cell viability and ISG induction was tested using quantitative reverse transcription PCR (RT-qPCR) of selected genes (Fig. S7A) and a cell viability assay (Fig. S7B).

**Fig. 7.**
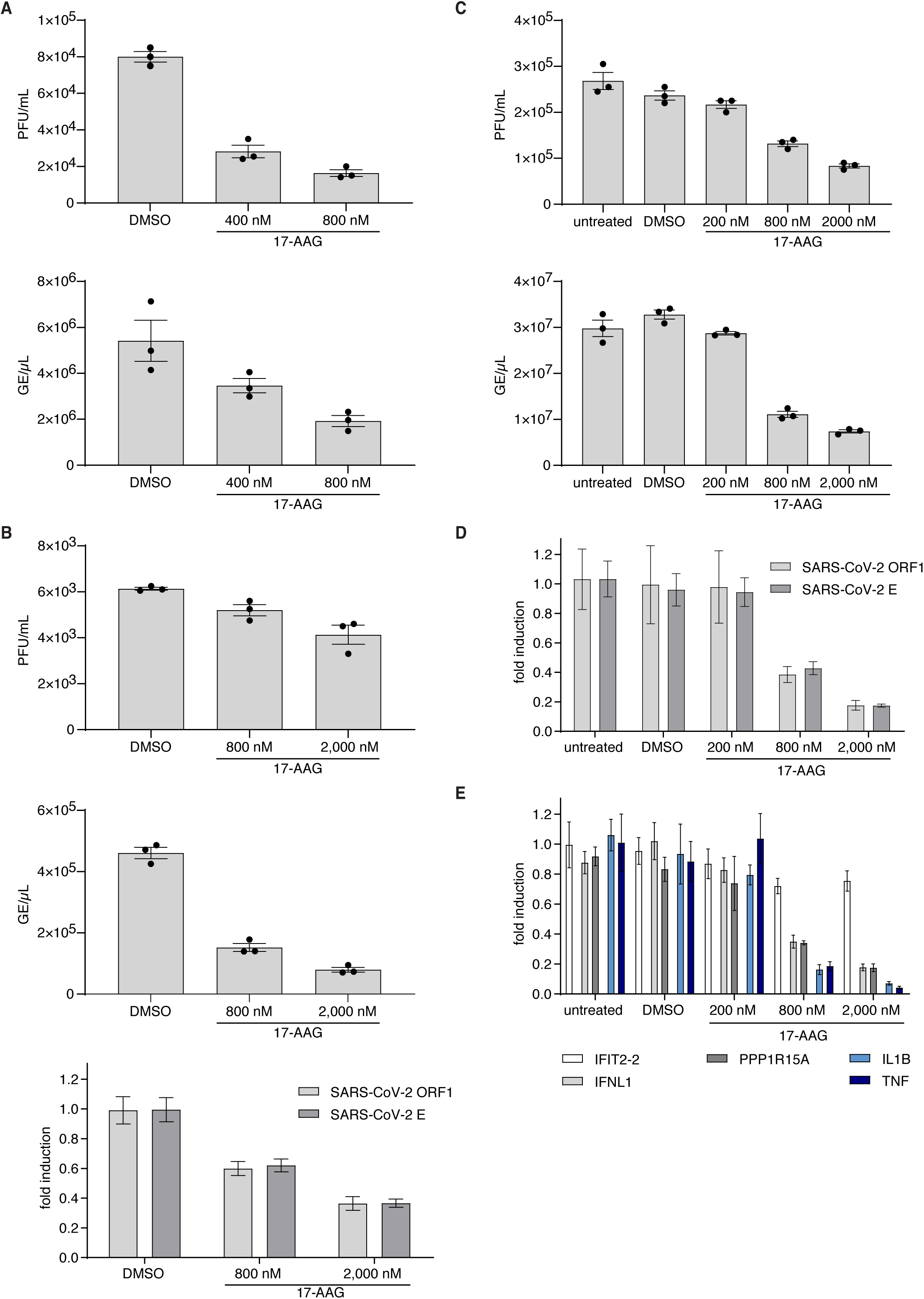
**A**, Infectious plaque forming units (PFU) (top) and viral RNA genome equivalents (GE) (bottom) in supernatants of Calu-3 cells infected with SARS-CoV-2 with the indicated treatment 16 hpi. After viral adsorption for one hour, cells were washed and supplied with conditioned medium containing DMSO, as solvent control, or indicated concentrations of 17-AAG. **B**, as in A but from an independent experiment analyzed at 8 hpi, and with additionally intracellular viral RNA. **C**, as in A, but with virus and 17-AAG added simultaneously without replacement of the inoculum. **D, E**, Intracellular expression of viral RNA and mRNA of selected genes from the samples in C. The 17-AAG treatment induces dose-dependent reduction of SARS-CoV-2.

## Discussion

We performed gene expression profiling of three different human cell lines infected with SARS-CoV and SARS-CoV-2 at bulk and single-cell level. We show a particularly strong induction of ISGs in Calu-3 cells, including cytokines, by SARS-CoV-2 in both bulk and scRNA-seq experiments. For various CoVs, a range of mechanisms that interfere with interferon signaling has been reported [60]. For SARS-CoV, it was shown that ORF6 inhibits signal transducer and activator of transcription (STAT) signaling [61] and that IRF3 activity is impaired [8]. Since RNA levels per cell (Fig. S4) and on the population level were comparable (Fig. 1), it is tempting to speculate that such mechanisms could be less efficient in SARS-CoV-2 compared to SARS-CoV. Indeed, cytokine production was described to be connected to pathogenesis [21, 40, 41]. However, the extent to which observations in cell lines and other models are reproducible in animal models and humans will require further investigations [62-64].

By comparing cell lines, we distinguish genes induced independently of the RNA sensing system, such as TXNIP and ARRDC3. Both genes are involved in signaling processes, and further investigations into their role in SARS-CoV-2 infection is warranted.

Small RNA profiling showed a strong induction of miR-155 in the infected cells. This miRNA has been associated with various virus infections [65-68]. miR-155-3p is also a well-known regulator of immune cells, in particular T-cell differentiation [69, 70]. Involvement of this miRNA in the regulation of innate immunity has also been reported [71]. Recently, miR-155-5p expression was shown to be induced in mice infected with influenza A virus [72]. Importantly, in this study, lung injury by ARDS was attenuated by deletion of miR-155, making this miRNA a potential therapeutic target in the context of COVID-19. Its role in SARS-CoV-2 infection and pathogenesis, whether it has a biological role in epithelial cells, and the potential for therapeutic interventions, e.g. through antisense oligonucleotide approaches, remains to be explored. The same holds true for the sRNAs generated from vtRNAs, and for other non-coding RNAs, which remain to be investigated in SARS-CoV-2 infection.

The scRNA-seq experiments provided a rich dataset to analyze host cell expression changes in response to infection. Surprisingly, the percentage of cells containing viral RNA was much higher than expected based on the MOI used for the infection experiments. This finding could also be explained by spreading of the cell-to-cell fusions, facilitated by the S protein on the cell surface [73]. Furthermore, the analysis of the scRNA-seq data of infected Calu-3 cells indicated a sequential activation of IRF and NF-κB target genes, and in particular, a putatively strong but transient induction of interferon genes. This could be due to a relatively short time window during the progression of infection, in which both IRF and NF-kB activity is sufficiently high to trigger interferon gene transcription [74, 75].

Concomitantly with the interferon induction, we observed a mild activation of ACE2 transcription in cells infected with SARS-CoV-2. Changes in ACE2 mRNA levels in the context of interferon treatment and coronavirus infection have been described before [58, 76]. Whereas the transcriptional induction (RNA velocity) was detectable, changes in mature mRNA levels were moderate in comparison to ISGs like IFIT1. Interestingly, SARS-CoV was shown to downregulate ACE2 protein levels in Vero cells [77].

Coronaviruses induce ER stress and activate the unfolded protein response (UPR) in infected cells [78, 79]. We observed transient induction of the stress-responsive heat shock protein gene HSP90AA1 [80], in the “slow-motion” infection model, H1299 cells, and 4hpi in Calu-3 cells. HSP90 modulates UPR by stabilizing the ER stress sensor transmembrane kinases IRE1α [81]. Inhibition of the HSP90 has previously been shown to slow down the replication of several viruses [59, 82, 83]. The reduction of SARS-CoV-2 growth by HSP90 inhibition was recently proposed based on a computational analysis of patient RNA sequencing data [84]. Here, we show that inhibition of HSP90 by 17-AAG at high nanomolar concentrations can reduce virus replication in an in vitro infection model. Interestingly, IFIT2 mRNA levels seemed unaffected by HSP90 inhibition, supporting the “on-off-switch” independent of the amount of viral RNA observed in the scRNA-seq data. In addition, mRNA expression of the pro-inflammatory cytokines TNF and IL1B, which are implied in the progression of COVID-19 [85], were strongly reduced. Since 17-AAG can induce apoptosis in cancer cells, which was not observed here for Calu-3 cells, the inhibitory effect of 17-AAG on viral replication needs to be expanded to studies in primary tissue infection models. Since several inhibitors of HSP90 with higher affinities have been in clinical development as anticancer agents [86] and advanced to phase 2 and 3 clinical trials, some of these compounds could be readily available to become part of a therapeutic strategy for COVID-19.

## Figure legends

Supplementary Fig. S1. A, Responsiveness to RNA as tested by RT-qPCR of three ISGs upon transfection of poly-I:C. B, growth kinetics of SARS-CoV and SARS-CoV-2 in the different cell lines (MOI 0.01). C, Coverage across the viral genome merged across all datasets for total RNA-seq, poly(A)+ RNA-seq, and small RNA-seq data. The top eight junctions supported by split reads are plotted in “sashimi” style for the total RNA-seq. D and E, Heatmaps of canonical junction-spanning reads, averaged across biological replicates per time point, expressed in TMM-normalized counts per million (D), or relative counts per time point (E). ORF1ab levels are estimated by counting contiguous reads mapping to the leader junction site. F, microscopy images (phase contrast) of infected Caco-2 and Calu-3 cells.

Supplementary Fig. S2. A, Log2 transformed fold changes of SARS-CoV infected cells at 12 hpi. Genes exhibiting, at least two-fold, significant changes in one of the two cell lines are shown in dark grey. Genes significantly deregulated in the same direction in both cell lines are shown in black, and labeled. All other genes are shown in light grey. B, Comparison of log2 transformed fold changes in Calu-3 in the two experiments from this study 12 hpi (series 1 and series 2). C-E, Comparison of log2 transformed fold changes in Calu-3 from series 2 of this study with various series of SARS-CoV-2 infected cells from Blanco-Melo et al. C, comparison with Calu-3 cells (series 7). D, Comparison with NHBE cells 12 hpi (series 1). E, comparison with A549 cells (series 2 and series 3 without, series 6 with ACE2 transduction).

Supplementary Fig. S3. A, Normalized counts (counts per million) of miR-155-3p (left panel) and miR-155-5p (right panel), colored by replicate. B, Log10 miRNA-155 host gene transcripts per million in samples measured by polyA- or total RNAseq. C, coverage plots of the three vaultRNA genes VTRNA1-2, VTRNA1-3, and VTRNA2-1.

Supplementary Figure S4. A, Percentages of cells bearing viral transcripts. B, Relative densities of the percentage of viral transcripts per cell (log10 transformed) for SARS-CoV (left) and SARS-CoV-2 infected cells (right). C, Cells are projected in two dimensions using UMAP and colored by time point, shaped by infection, and split by replicates. D, Two-dimensional projection of cells colored by cluster. E, For each sample, the number of cells per cluster was divided by the total number of cells from this sample. The contributions of these values to the composition of the clusters is displayed by a stacked bar plot. F, Heatmap of expression levels of cluster markers. G, H, J, M, As in C but colored by expression values of the indicated genes. I, Distribution of expression of the miR-155 host gene MIR155HG per cluster. K, Correlation of gene expression values with the amount of SARS-CoV RNA (x-axis) and SARS-CoV-2 RNA (y-axis). L, Correlation of gene expression values with the amount of IFNB1 (x-axis) and ARRDC3 (y-axis). N, Clustering of cells without viral transcripts, analogous to figure 4 and S4C.

Supplementary Figure S5.

A-C, Projection as in Fig. 5A, colored by the indicated expression values. D, Heatmap with intron TPM counts for selected genes, columns depict cells and are ordered by ascending SARS-CoV-2 load per cluster. E, Upper panel as in A-C, lower panel barplot showing fraction of cells containing intron reads from the ACE2 gene.

Supplementary Figure S6. A, Shows percentages of cells with virus. B-D, H1299 cells are projected in two dimensions using Uniform Manifold Approximation and Projection (UMAP) and colored as indicated.

Supplementary Figure S7. The cytotoxicity of the HSP90 inhibitor was measured by RT-PCR of selected genes (A) and a cell viability assay (B). The activity of untreated cells was set as 100% and the cell viability with the indicated treatment is shown as fold induction over untreated cells after 16 and 24 hpi.

## Materials and Methods

### Cell culture

Vero E6 (ATCC CRL-1586), Calu-3 (ATCC HTB-55), Calu-2 (ATCC HTB-37) and H1299 (ATCC CRL-5803) were cultivated in Dulbecco’s modified Eagle’s medium (DMEM) supplemented with 10% heat-inactivated fetal calf serum, 1% non-essential amino acids, 1% L-glutamine and 1% sodium pyruvate (all Thermo Fisher Scientific) in a 5% CO_2_ atmosphere at 37 °C.

### Poly-I:C transfections

Transient transfection of eukaryotic cells was performed using X-tremeGENE™ siRNA transfection reagent (Roche) according to the manufacturer’s instructions. Briefly, 2×10^5 cells/ml were grown in 6-well plates for 24 h and fresh DMEM without antibiotics was added. OptiPRO SFM™ (Gibco) was supplemented with 0.25 µg poly(I:C) (Invivogen) and 0.75 µl X-tremeGENE™ siRNA reagent, incubated for 15 min, and 100 µl transfection mix was added to the cells.

### RT-qPCR on intracellular RNA

RNA was isolated from Trizol using the RNA clean and concentrator kit (Zymo). The RNA was reverse transcribed using maxima RT and subjected to qPCR as described [26]. Primers used for qPCR are listed in supplementary table 5.

### Viruses

SARS-CoV (Frankfurt strain, NCBI accession number AY310120) and SARS-CoV-2 (Patient isolate 985, BetaCoV/Munich/BavPat1/2020|EPI_ISL_406862) were used. For virus stock production, virus was grown on Vero E6 cells and concentrated using Vivaspin® 20 concentrators (Sartorius Stedim Biotech). Virus stocks were stored at −80°C, diluted in OptiPro serum-free medium supplemented with 0.5% gelatine (Sigma Aldrich) and phosphate-bufferd saline (PBS, Thermo Fisher Scientific). Titer was defined by plaque titration assay. Cells inoculated with cell culture supernatants from uninfected Vero cells mixed with OptiPro serum-free medium supplemented with 0.5% gelatine and PBS, in accordance to virus stock preparation, serves as mock infected controls. All infection experiments were carried out under biosafety level three conditions with enhanced respiratory personal protection equipment.

### Virus growth kinetics and plaque titration assay

24 h prior to infection, the different cell lines were seeded to 70% confluence. The cells were washed once with PBS before virus (diluted in OptiPro serum-free medium) adsorption. After incubation for 1 h at 37 °C, 5% CO2 the virus-containing supernatant was discarded and cells were washed twice with PBS and supplied with DMEM as described above.

To determine the amount of infectious virus particles in the supernatant a plaque titration assay was performed. For the assay Vero E6 cells were seeded to confluence and infected with serial dilution of virus-containing cell culture supernatant diluted in OptiPro serum-free medium. One hour after adsorption, supernatants were removed and cells overlaid with 2.4% Avicel (FMC BioPolymers) mixed 1:1 in 2xDMEM. Three days post-infection the overlay was removed, and cells were fixed in 6% formaldehyde and stained with a 0.2% crystal violet, 2% ethanol and 10% formaldehyde.

### Western Blot Analysis

The expression of human ACE-2 (hACE-2) was confirmed by Western blot analysis. For the preparation of total cell lysate cells were washed with PBS and lysed in RIPA Lysis Buffer (Thermo Fisher Scientific) supplied with 1% Protease Inhibitor Cocktail Set III (Merck Chemicals). After an incubation of 30 min at 4 °C, cell debris were pelleted (10 min, 13,000 x g, 4 °C) and the supernatant transferred to a fresh reaction tube. For determining protein concentration Thermo Scientific’s Pierce™ BCA Protein Assay Kit, according to the manufacturer’s instructions was used. The protein lysates were mixed with 4xNuPAGE LDS Sample Buffer (Invitrogen) supplemented with 10% 2-mercaptoethanol (Roth). Protein lysates were separated by size on a 12% sodium dodecyl sulfatepolyacrylamid (SDS) gel and blotted onto a 0.2 µm polyvinylidene difluoride (PVDF) membrane (Thermo Scientific) by semi-dry blotting (BioRad). Primary detection of hACE-2 was done using a goat anti-hACE-2 antibody (1:1,250; #AF933, R&D Systems), a horseradish peroxidase (HRP)-labeled donkey anti-goat antibody (1:5,000, Dianova) and Super Signal West Femto Chemiluminescence Substrate (Thermo Fisher Scientific). As loading control, samples were analyzed for β-actin expression using a mouse anti-β-actin antibody (1:5,000, Sigma Aldrich) and a HRP-labeled goat anti-mouse antibody (1:10,000, Sigma-Aldrich).

### Infections for RNA sequencing experiments

Calu-3 cells and H1299 cells were seeded at a concentration of 6 x 10^5 cells/mL and 5 x 10^4 cells/mL, respectively. 24 h post seeding cells were infected with SARS-CoV and SARS-CoV-2 at an MOI of 0.33 or Vero cell supernatant mixed with Optipro serum-free medium supplemented with 0.5 % gelatine and PBS as negative control. 4, 8, 12 and 24 hpi samples were taken. For sequencing of total RNA the supernatant was removed and Trizol LS Reagent (Thermo Fisher Scientific) was applied to the cell-layer and incubated for 1 min at room temperature until the cells were lysed. The suspension was then transferred to a RNase free reaction tube (Thermo Fisher Scientific) and stored at −80 °C.For scRNA-seq sample preparation the cells were washed with pre-warmed PBS, detached with pre-warmed trypsin for 3 min at 37 °C. The detached cells were transferred into a reaction tube (Eppendorf) and the following steps were performed on ice. Cells were spinned down at 1000 x g for 2 min at 4 °C, resuspended in PBS properly and passed through a 35 µm blue snap cap cell strainer (STEMCELL) and again pelletized The cell pellet was then resuspended in pre-chilled methanol (Roth) and stored at −80 °C.

### RNA sequencing

#### Poly-A RNA sequencing

Poly-A RNA sequencing libraries were prepared using the NEBNext Ultra II Directional RNA Library Prep Kit (NEB) according to the manufacturer’s protocols. Libraries were sequenced on a NextSeq 500 device at 1×76 cycles.

#### Small RNA sequencing

100 ng of total RNA of each condition was used for small RNA library preparation. Library preparation was performed using the SMARTer smRNA-Seq kit for Illumina from Clontech according to manufacturer’s instruction. The small RNA libraries were pooled together with 19 % PhiX and sequenced on the NextSeq 500, 1 × 50 cycles.

#### Total RNA sequencing

1 µg of total RNA of each condition was used for total RNA library preparation. First, samples were depleted of ribosomal RNA using the RiboCop rRNA Depletion Kit (Lexogen) according to manufacturer’s instruction. Following, ribo-depleted samples were processed with the TruSeq mRNA stranded kit from Illumina according to manufacturer’s instruction. The total RNA libraries were sequenced on the HiSeq 4000, 2 x 76 cycles.

#### Viral RNA-seq analysis

Total and poly(A)^+^ RNA-seq reads were mapped with STAR 2.7.3a to a combined genome comprised of GRCh38 and GenBank MN908947 (SARS-CoV-2) or AY310120 (SARS-CoV) using permissive parameters for noncanonical splicing [33, 91]. Viral genes were quantified by taking the top eight noncanonical splice events called by STAR across all total RNA-seq datasets according to the numbers of uniquely-mapping reads spanning the junction (Supplemental Table 3). To estimate levels of ORF1ab, insertions, soft-clipping events and split reads were filtered from virus-mapping reads, followed by intersection with positions 53-83 of the virus using bedtools, requiring a minimum of 24 nucleotides overlap to reflect the parameters STAR requires to call a noncanonical splice junction [92]. These counts were either combined with a count matrix of the human genes quantified by STAR and TMM/CPM normalized with edgeR (Figure S1D) or normalized by the total number of viral junction-spanning reads per time point (Figure S1E) [93]. Coverage plots were made from merged STAR-mapped BAM files, or from Bowtie-mapped small RNA-seq BAM files using ggsashimi [94]. This workflow was implemented with custom Python scripts in a Snakemake pipeline [95].

#### microRNA analysis

Raw reads were preprocessed by trimming with cutadapt (version 2.9) in two passes, first trimming i) the Illumina TruSeq adaptor at the 3’ end and allowing for one mismatch, ii) all 3’end bases with mean Phred score below 30 and iii) the three 5’end overhang nucleotides associated with the template-switching Clontech library preparation protocol.

In the second pass, poly(A)-tails were trimmed. Trimmed reads were mapped using bowtie (version 1.2.2) to a SARS genome consisting of the combined SARS-CoV and SARS-CoV-2 genomes using the non-standard parameters (-q -n 1 -e 80 -l 18 -a -m 5 –best – strata). Reads that did not align to the SARS-CoV genome were aligned to the GRCh38 genome. The expression of known miRNAs (miRBase 22 annotation) was estimated using mirdeep2 (version 2.0.0.7) and standard parameters.

The differential expression analysis used the limma [96] and edgeR [93] packages after applying the voom transformation to the TMM-normalized count data produced by mirdeep2. For the different viral infections we contrasted SARS-CoV2-24h – SARS-CoV-2-4h with mock-24h - mock-4h in order to test for those miRNAs differentially expressed long after the infection having removed any effects seen in mock as well.

#### Bulk RNA-sequencing analysis using DESeq2

Starting from count tables, RNA sequencing results were analysed on a per run basis comparing infected samples to time matched mock experiments unless otherwise specified using DESeq2 [97] version 1.22.2. Genes with a maximum read count across samples of less than two were filtered out. Differentially expressed genes were defined as genes with an absolute fold change in mRNA abundance greater than 1.5 (log2 fold change of 0.58 - using DESeq2 shrunken log2 fold changes) and an adjusted p-value of less than 0.05 (Benjamini-Hochberg corrected).

#### Gene ontology and KEGG enrichment analysis

Genes whose mRNAs were found to be differentially expressed were subjected to gene set overrepresentation analysis using the clusterProfiler package in R [98]. Specifically, gene sets from Gene Ontology (Molecular Function, Biological Process, Cellular Component) and KEGG pathways containing

#### Single-cell RNA-seq

Methanol-fixed cells were centrifuged at 2,000 x g for 5 min, rehydrated in 1 mL rehydration buffer containing 0.01% PBS/BSA and 1:100 Superasein (Thermo Fisher), and resuspended in 400 µL rehydration buffer followed by passing through a 40 µm cell strainer. Encapsulation was done on the Nadia system (Dolomite biosystems) using the built-in standard procedure. For library preparation, we followed the version 1.8 of the manufacturer’s protocol, with adding a second-strand synthesis step [99].

For the encapsulation, 75,000 cells in 250 µL rehydration buffer were used, with 250 µL of lysis buffer (6% Ficoll PM-400, 0.2% Sarkosyl, 20 mM EDTA, 200 mM Tris pH 7.5, 50 mM DTT) and 3 mL oil (Biorad #1864006). After encapsulation, beads were recovered from the emulsion by washing with 2 × 30 mL 6 × saline sodium citrate buffer (diluted from Sigma #S6639) buffer in a 5 µm ÜberStrainer (pluriSelect). After another washing step in 1.5 mL 6 x SSC, cells were washed with 5 x reverse transcription (250 mM Tris pH 8, 375 mM KCl, 15 mM MgCl2, 50 mM DTT) buffer and resuspended in 200 µL RT mix (50 mM Tris pH 8, 75 mM MgCl2, 3 mM MgCl2, 10 mM DTT, 4% Ficoll PM-400, 1 mM each dNTPs, 2.5 µM Macosko TSO, 10 µl Maxima H-RT enzyme). Beads were incubated for 30 min at room temperature and 90 min at 42 °C (all incubation steps on a rotator). After washing once with TE/0.5% SDS and twice with TE/0.01% Tween, beads were incubated in 200 µL exonuclease mix (10µl Exonuclease in 1xexonuclease buffer, NEB #M0293) for 45 min at 37 °C, again on a rotator. After washing with once with TE/0.5% SDS and twice TE/0.01% Tween, beads were incubated for 5 min in 0.1 M NaOH, washed with TE/0.01% Tween and TE, and incubated in 200 µl second strand mix (50 mM Tris pH 8, 75 mM MgCl2, 3 mM MgCl2, 10 mM DTT, 12% PEG 8000, 1 mM each dNTPs, 10 µM dN-SMRT oligo, 5 µl Klenow enzyme NEB #M0212) for 1 h at 37 °C. Beads were again washed in TE/0.01% Tween and stored overnight in TE/0.01% Tween, then washed in TE and twice in water, and per sample eight PCR reactions with 4,000 beads each in 50µl using 1µM SMART PCR primer (oligos in supplementary table 5) and the 2x Kapa HiFi Hotstart Ready mix (Roche #07958935001) were performed, with pre-incubation for 3 minutes at 95 °C, then 4 cycles 98 °C/20s, 65°C/45s, 72 °C/3min, then 9 cycles 98 °C/20s, 67°C/20s, 72 °C/3min, then post-incubation for 3 minutes at 72 °C. The eight PCR reaction were pooled in three clean-up reactions using Ampure XP beads. For each oft the three sub-samples, a Nextera XT v2 (Illumina) reaction was performed with 600 pg DNA. In a 20µl reaction, 10 µl tagment DNA buffer and 5 µl amplicon tagment mix were incubated for 5 minutes at 55 °C, and, after addition of 5 µl neutralization buffer for 5 minutes at room temperature. Afterwards, 15 µl PCR master mix were added, 200 nM New-P5-SMART PCR hybrid oligo, 200 nM index oligo in total 50 µl. The Nextera reactions were then again pooled, purified using Ampure XP beads, and sequenced on a NovaSeq 6000 deviced with 21+71 cycles using Read1CustomSeqB for read 1.

#### Single-cell data processing

After trimming one nucleotide from the 3’ end of read one, count tables were generated using the pigx scRNA-seq pipeline [100] version 1.1.4.

#### Single-cell data analysis

All analysis shown in figure 4-6 was done similar as described previously [26] using Seurat and ggplot2 packages [101, 102].

#### HSP90 inhibitor experiments

The HSP90 inhibitor 17-AAG was purchased from Sigma () and dissolved in DMSO. Cells were seeded and grown to subconfluence and infected with SARS-CoV-2 MOI 0.01 diluted in OptiPro serum free medium. After 1 h virus adsorption the supernatant was removed and cells were washed twice with PBS. DMEM containing dilutions of 17-AAG (200 nM, 400 nM, 800 nM, 2,000 nM) or DMSO as solving control. Samples for detection of viral RNA and infectious particles in the supernatant as well as total RNA within the cells were taken 8, 16 and 24 hpi.

The cytotoxicity of the the HSP90 inhibitor was assured by cell viability assay using CellTiter-Glo® Luminescent Cell Viability Assay according to manufacturer’s instruction (Promega). The activity of untreated cells was set as 100% and cells were treated with different concentrations of 17-AAG. The viability of cells was measured 16 and 24 h after treatment using Mithras Luminescence microplate reader (Berthold).

#### RT-qPCR of viral RNA in the supernatant

The viral RNA from supernatant of infected cells was isolated using the NucleoSpin RNA virus isolation kit (Macherey-Nagel) according to the manufacturer’s instructions. To determine the amount of viral genome equivalents the previously published assay specific for both SARS-CoV and SARS-CoV-2 Envelope gene [103] was used. Data analysis was done using LightCycler Software 4.1 (Roche).

## Data availability

Raw sequencing is available at the Gene Expression Omnibus database (GEO), identifier GSE148729 (https://www.ncbi.nlm.nih.gov/geo/query/acc.cgi?acc=GSE148729). Supplementary data and supporting files such as scRNA-seq Seurat objects are available at www.mdc-berlin.de/singlecell-SARSCoV2.

## Acknowledgements

The authors wish to thank Melanie Brinkmann, Leif Sander, Marco Hein, Joseph Luna, Friedemann Weber and Robert Zinzen for comments and discussion; Jeannine Wilde, Tatiana Borodina, Daniele Franze and Nouhad Benlasfer for sequencing and technical assistance; and members of the Landthaler/Drosten/Rajewsky/Akalin/Selbach labs for continuous support under extraordinary circumstances.

## Contributions

E.W. performed experiments, analyzed data, and wrote the manuscript, K.M. performed infection and inhibitor experiments, V.F. performed coding and data analysis, A.D. performed smallRNA and total RNA sequencing together with S.A., L.T.G performed infection experiments and contributed to manuscript writing, R.A. performed experiments, F.K. analyzed smallRNA and totalRNA sequencing data together with I.L. and A.I., D.K. analyzed viral transcripts, C.B. analyzed gene expression together with T.M., S.D.G. and J.P.P. performed experiments, D.N., M.A.M., M.A., A.A., N.R. supervised parts of the project, and C.D. and M.L. were responsible for overall supervision.

