## Supplementary Figures for "Bulk and single-cell gene expression profiling of SARS-CoV-2 infected human cell lines identifies molecular targets for therapeutic intervention"

Supplementary figure S1, part 1

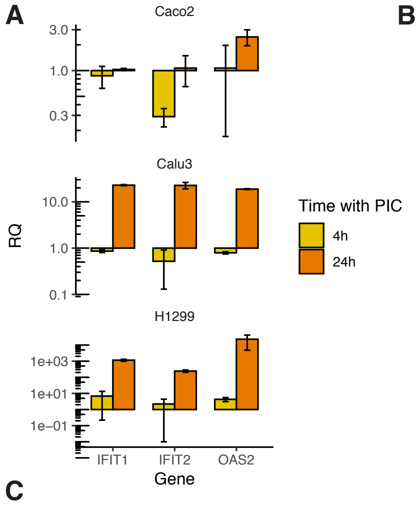


B

All samples: SARS-CoV and SARS-CoV-2, duplicates

| | Calu-3 epthelial lung adenocarcinoma poly-I:C responsive ACE2 high, TMPRSS2 ok | | | | | H1299 epithelial lung adenocarcinoma poly-I:C responsive ACE2 low, no TMPRSS2 | | Caco-2 epithelial colorectal adenocarcinoma Not poly-I:C responsive ACE2 low, TMPRSS2 ok |
|---|---|---|---|---|---|---|---|---|
| | polyA RNA 1 | polyA RNA 2 | single-cell RNA | small RNA | total RNA | polyA RNA | single-cell RNA | polyA RNA |
| 4 h | + | + | + | + | + | + | + | + |
| 8 h | | + | + | | | | | |
| 12 h | + | + | + | + | + | + | + | + |
| 24 h | + | | | + | + | + | + | + |
| 36 h | | | | | | + | | |

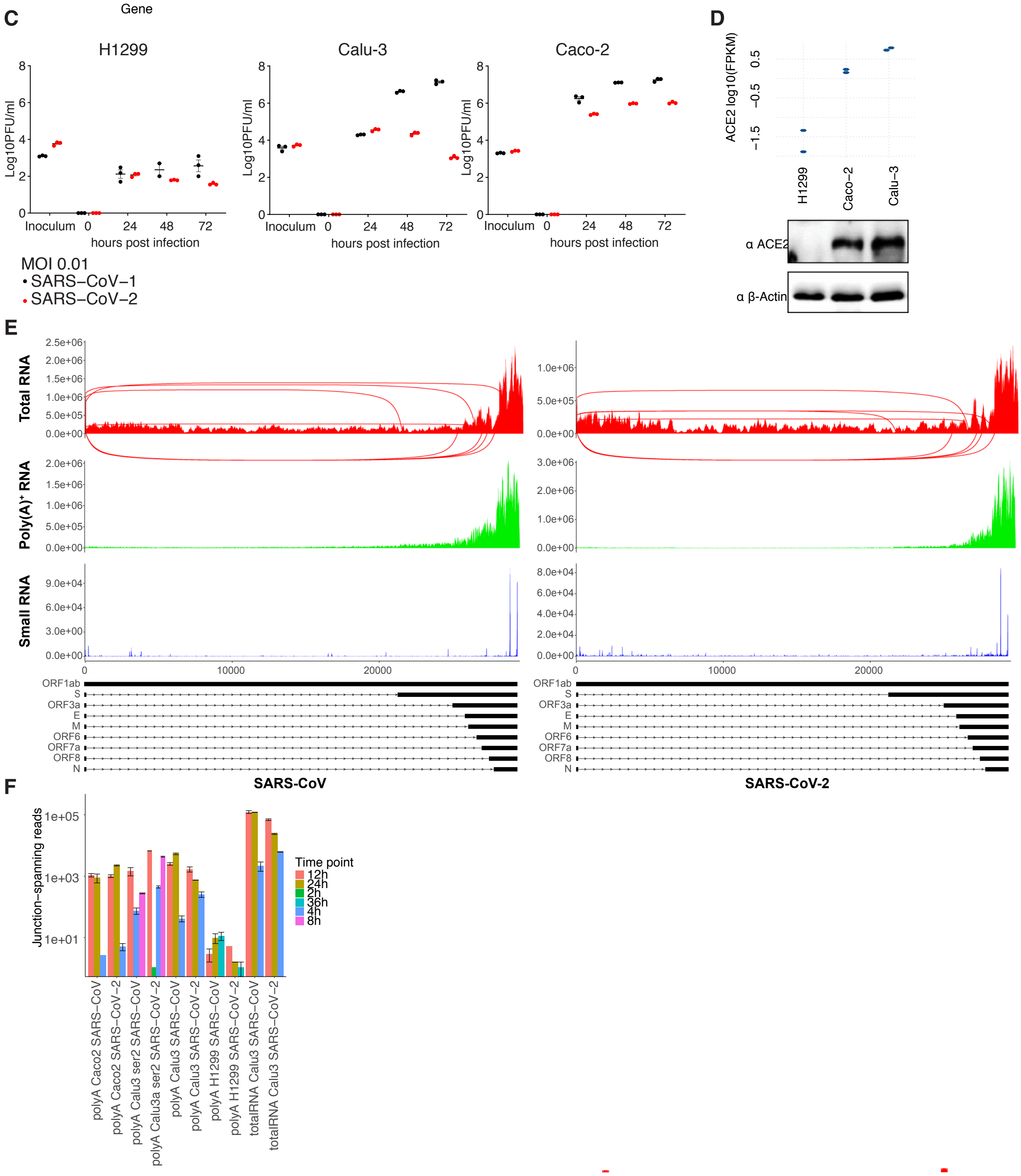

Supplementary Figure S1, part 2

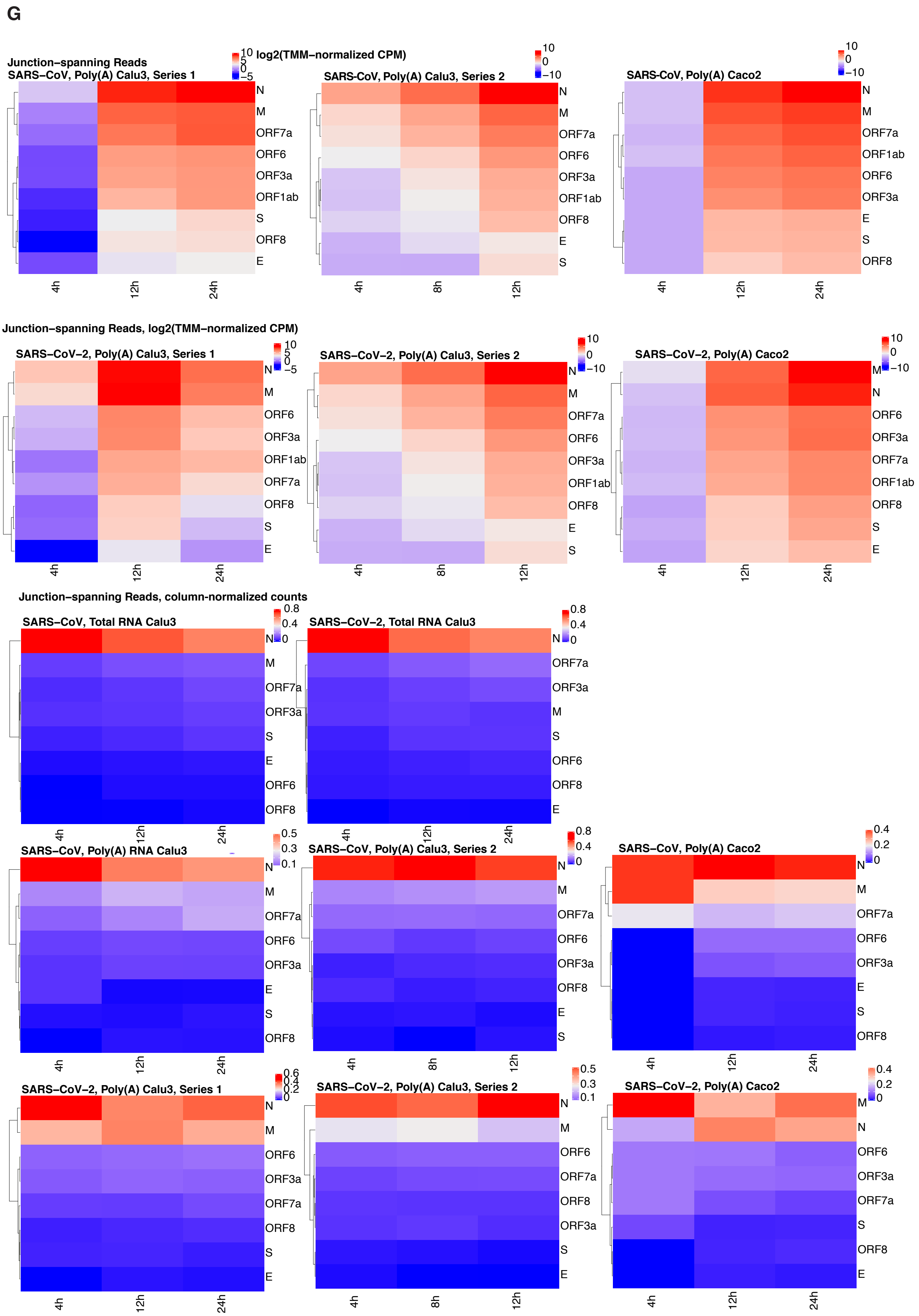

Supplementary Figure S1, part 2

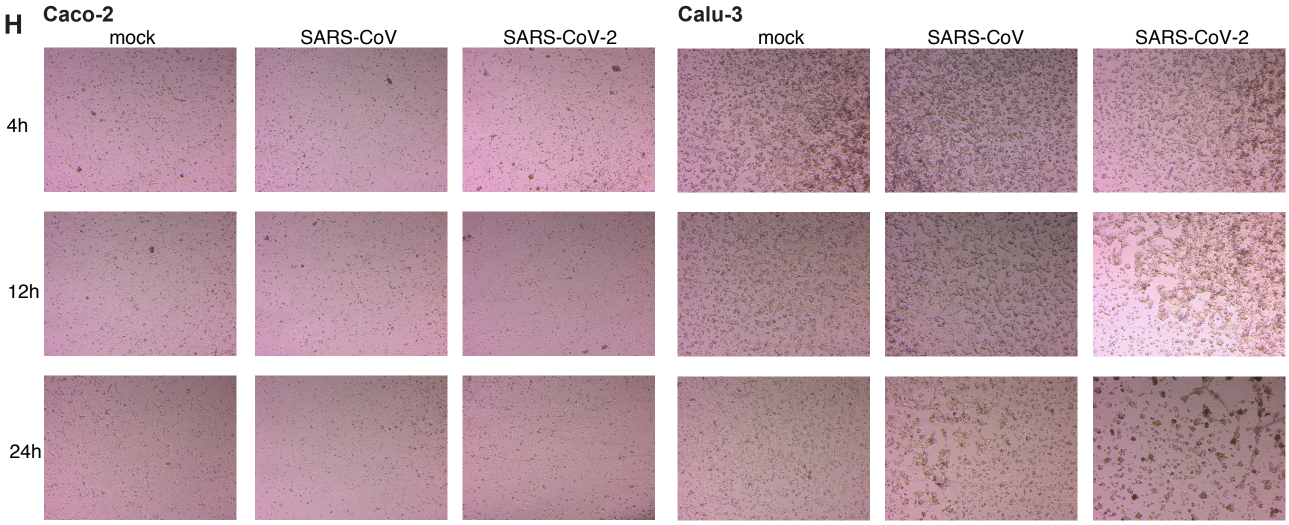

Supplementary Figure S2, part 1

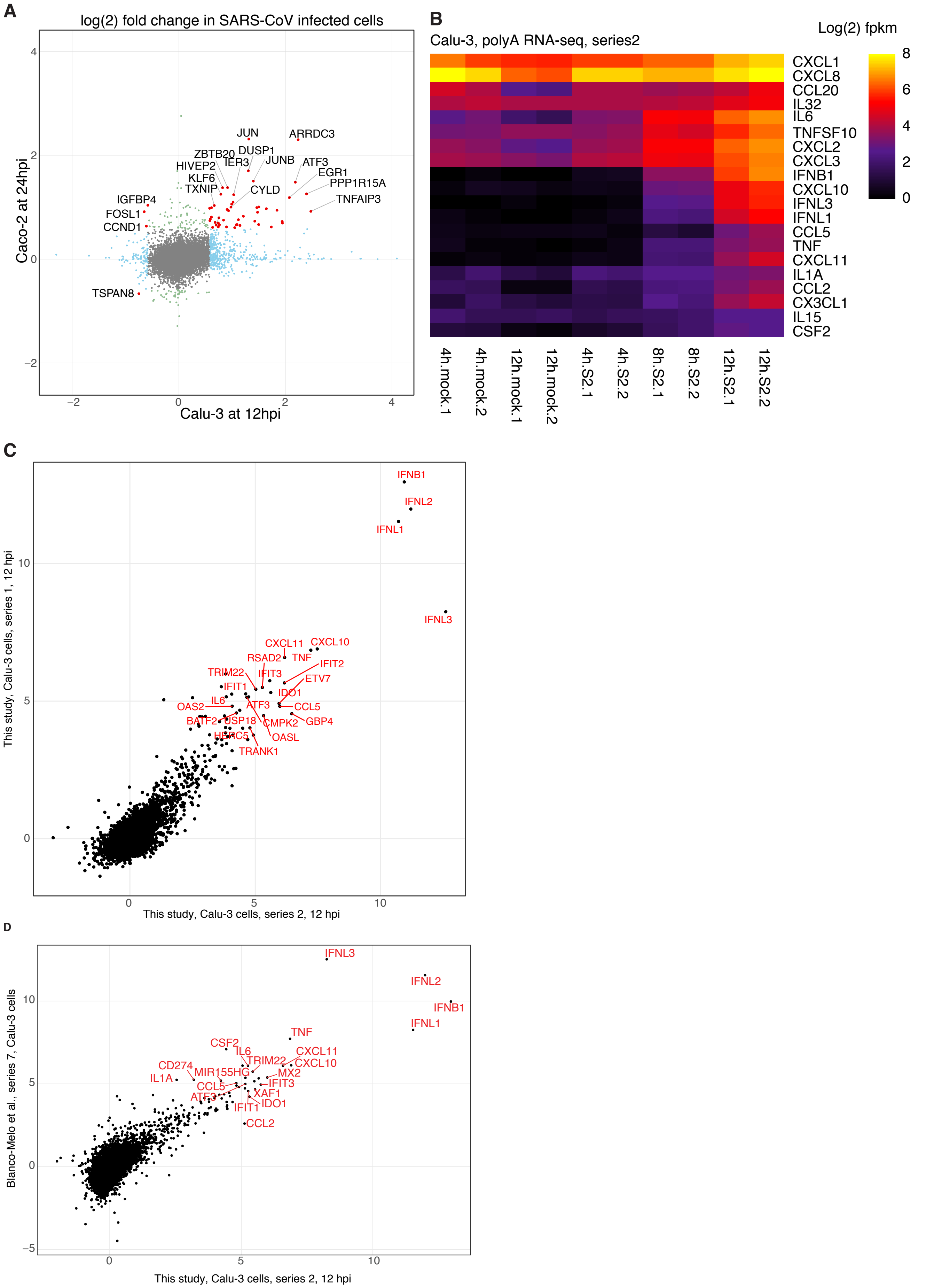

Supplementary Figure S2, part 2

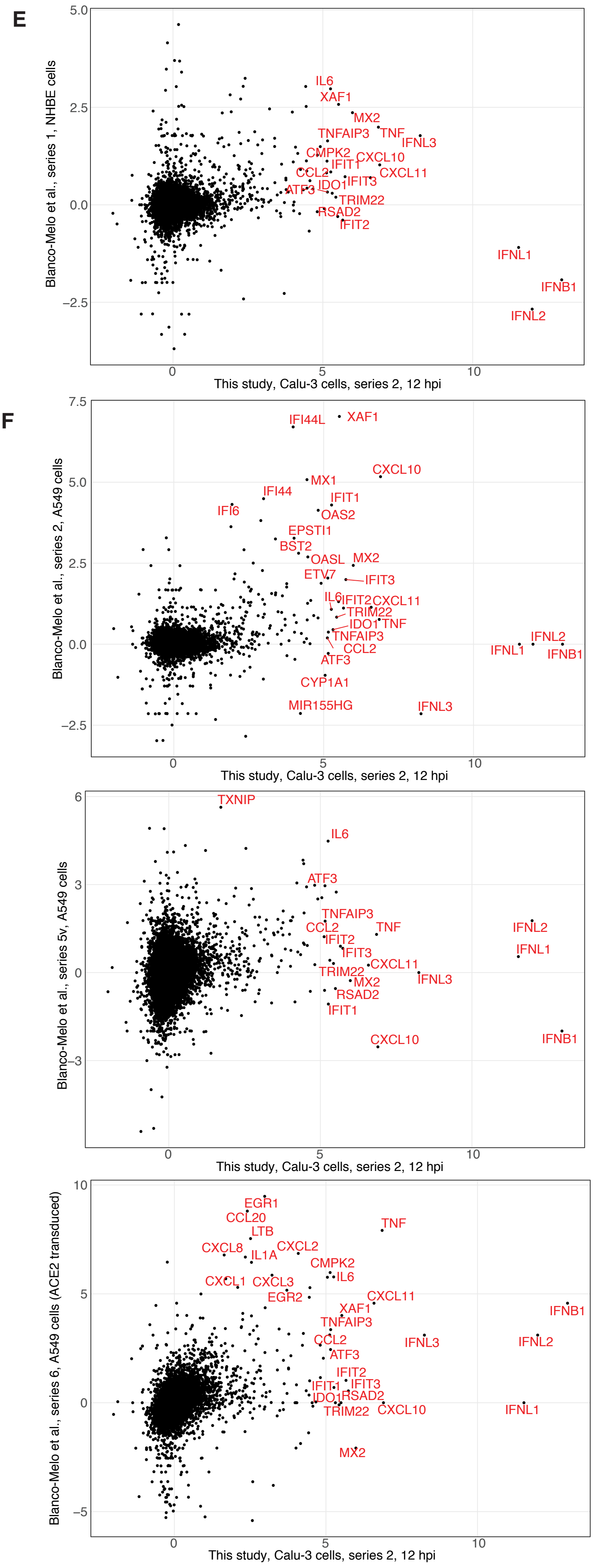

Supplementary Figure S3

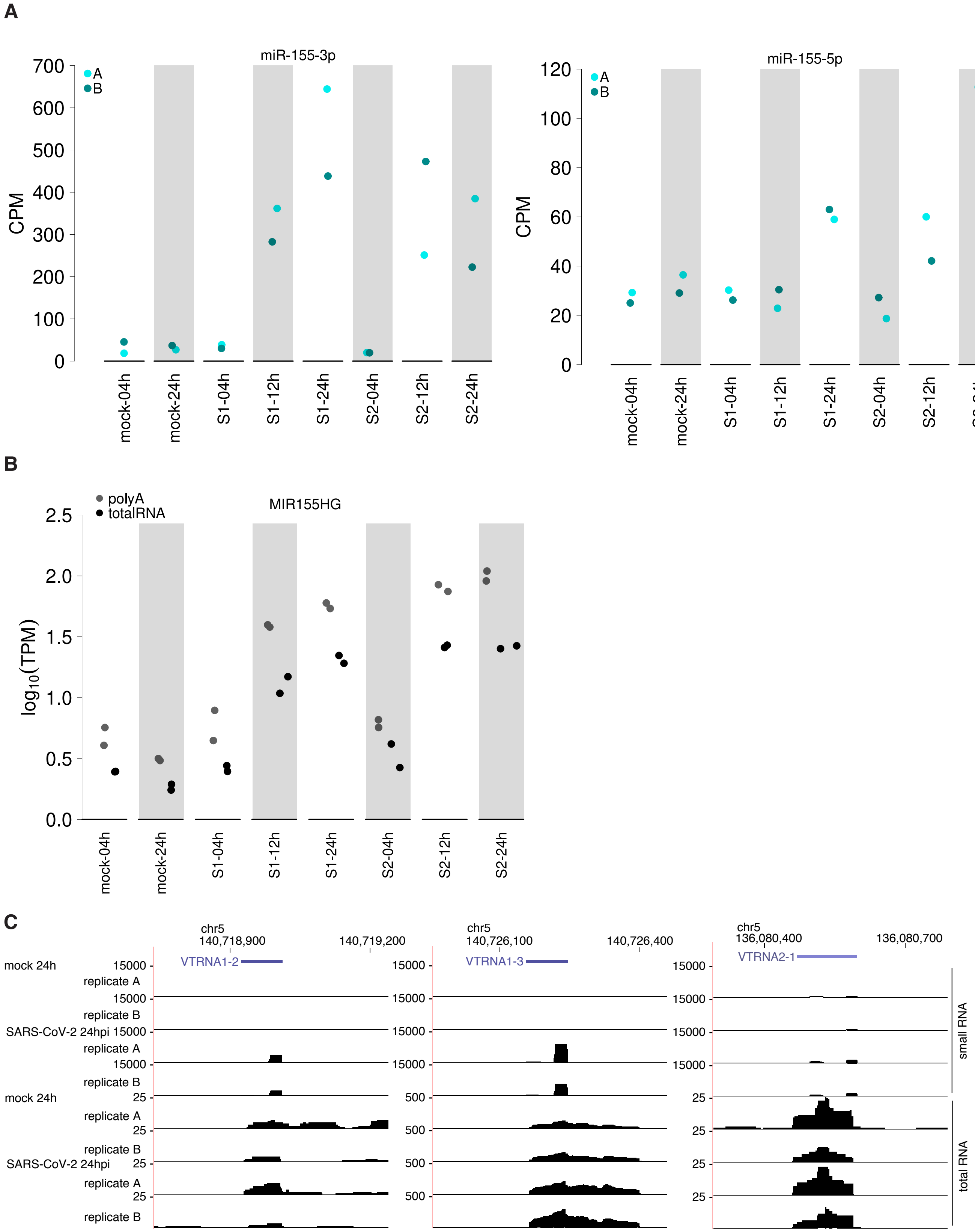

Figure S4, part 1

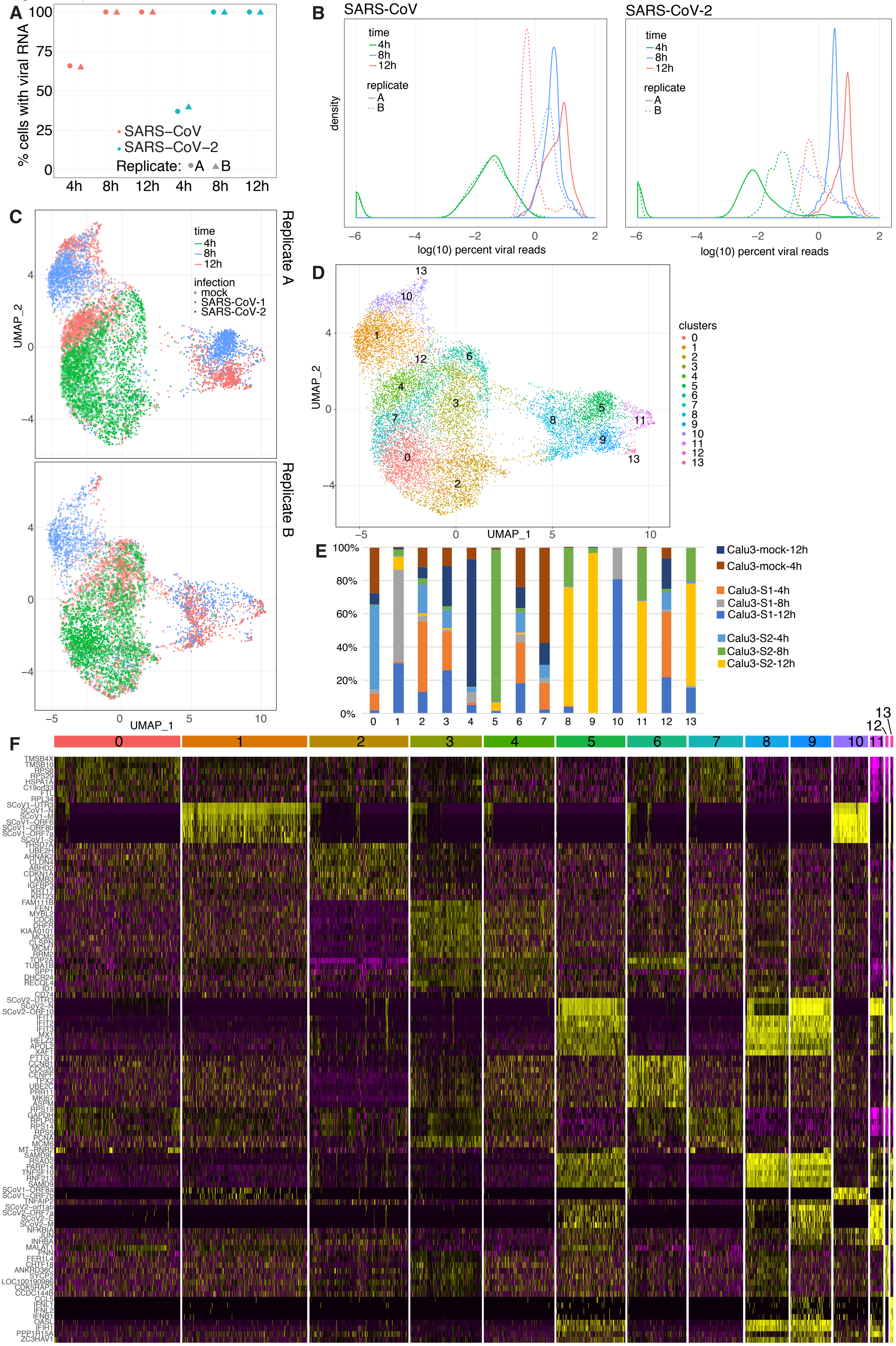

Figure S4, part 2

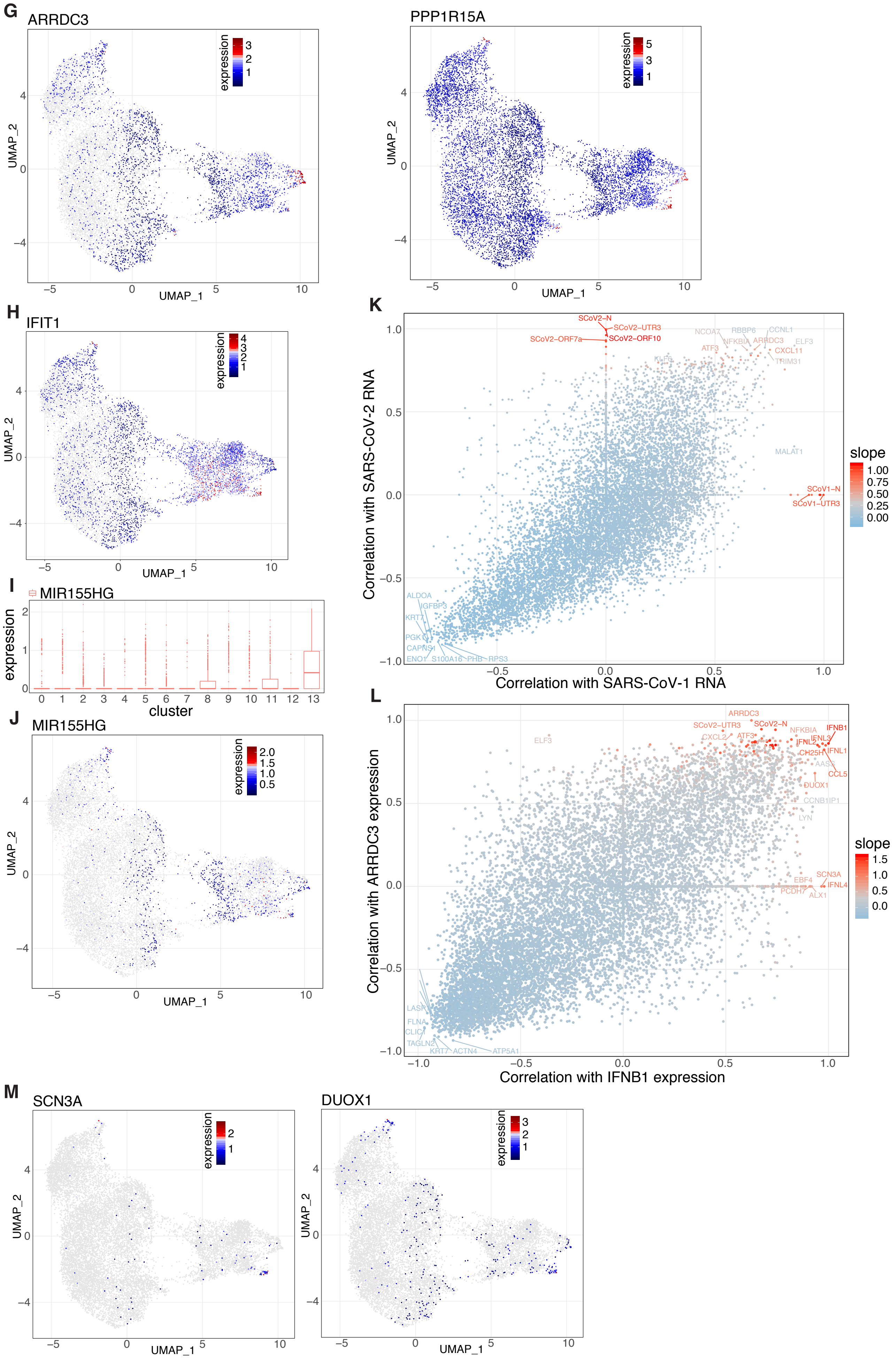

Figure S4, part 3

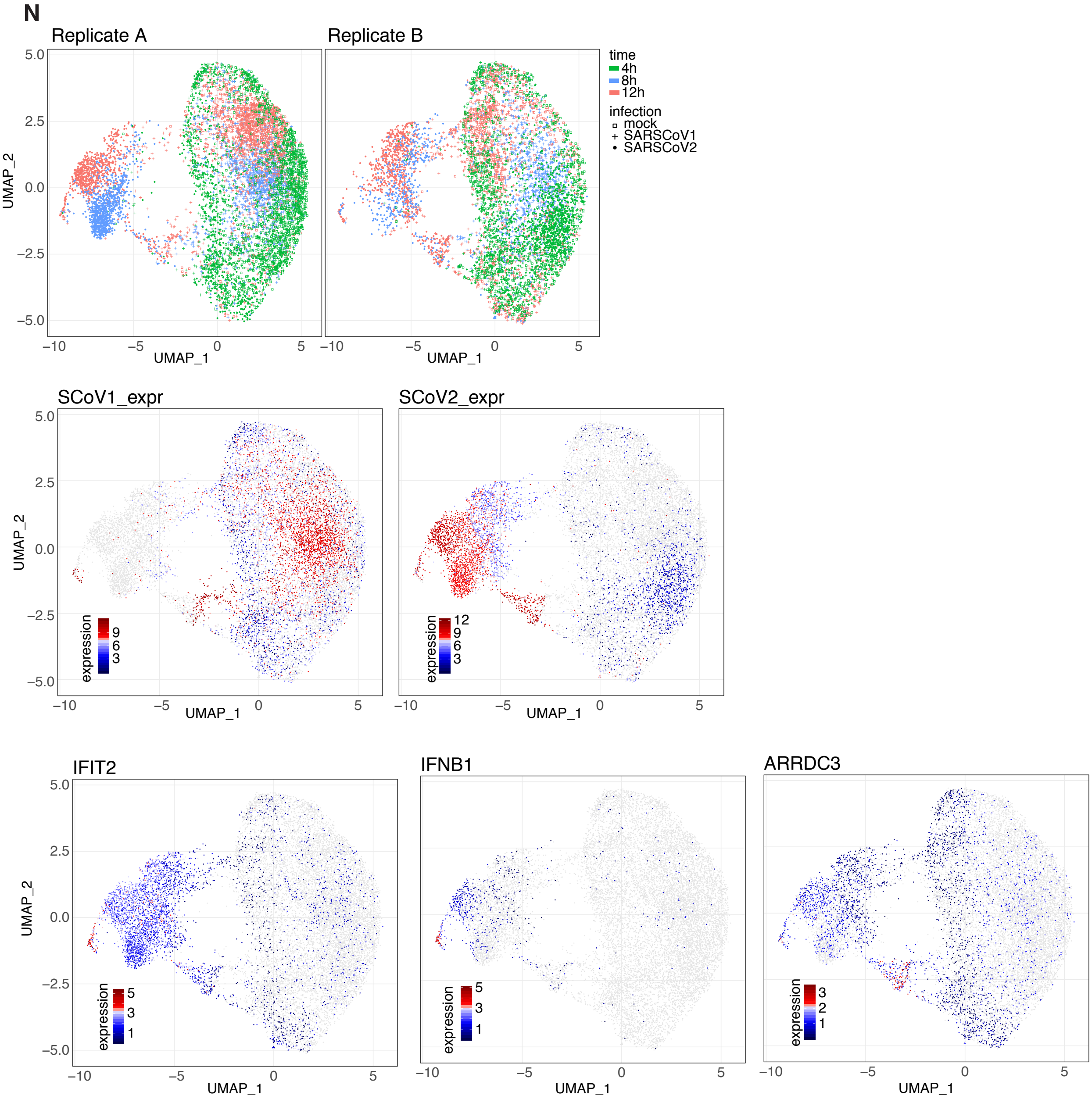

Figure S5

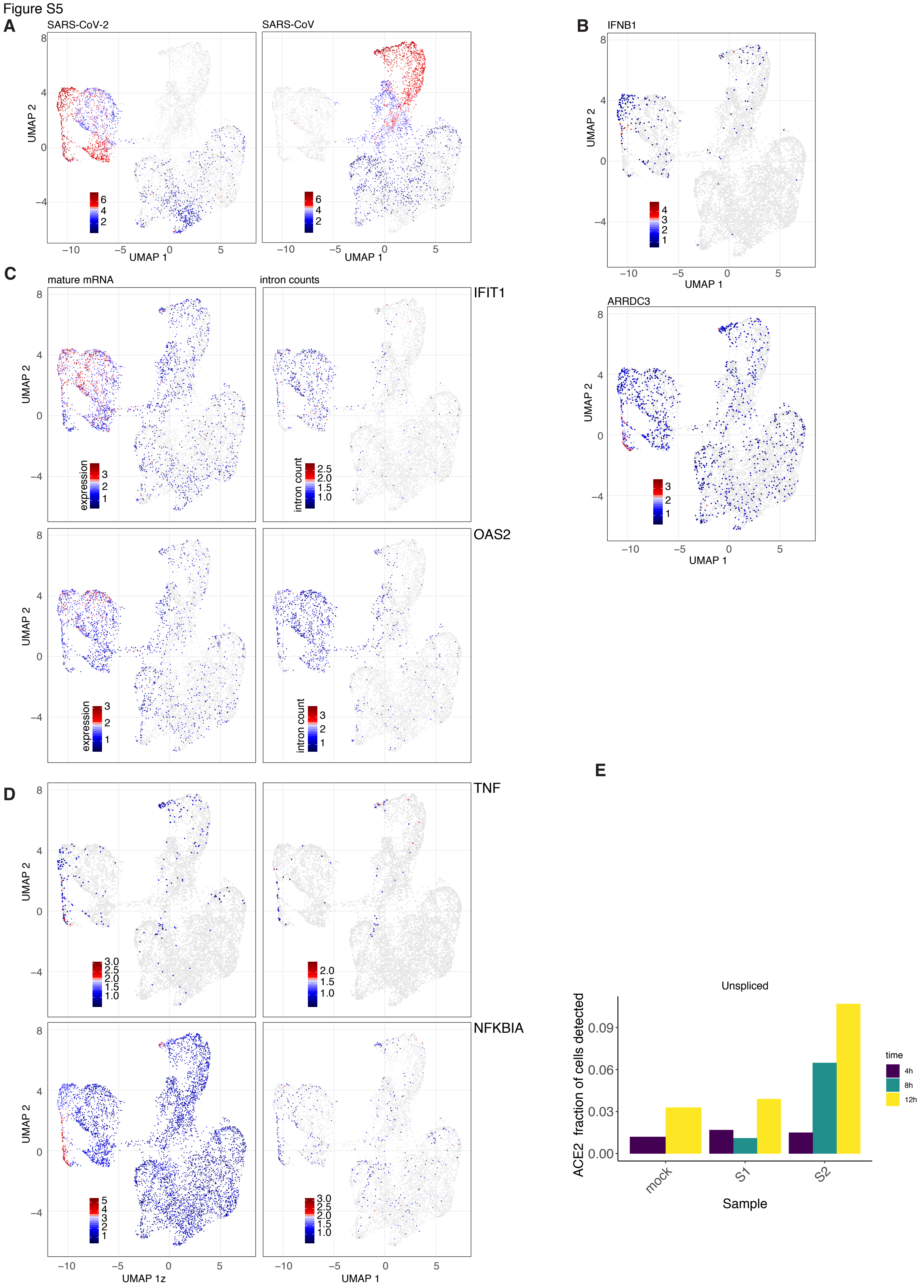
A
SARS-CoV-2
SARS-CoV
UMAP 2
UMAP 1
B
IFNB1
ARRDC3
C
mature mRNA
intron counts
IFIT1
OAS2
expression
intron count
D
TNF
NFKBIA
UMAP 1z
E
Unspliced
ACE2 fraction of cells detected
Sample
mock
S1
S2
time
4h
8h
12h

Figure S5

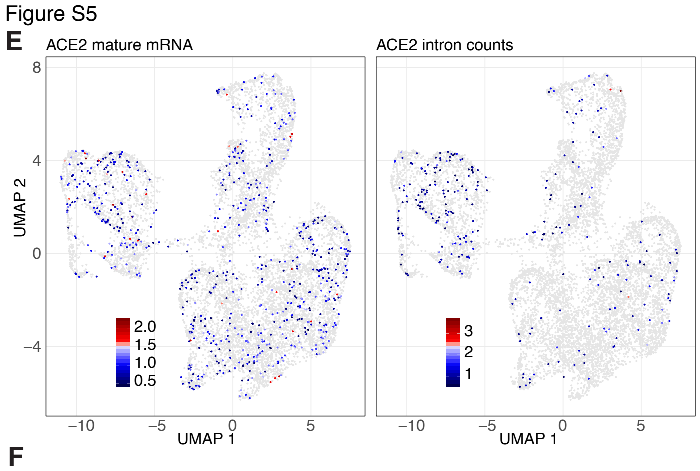

E
ACE2 mature mRNA
ACE2 intron counts
UMAP 2
UMAP 1
8
4
0
−4
−10
−5
0
5
2.0
1.5
1.0
0.5
3
2
1


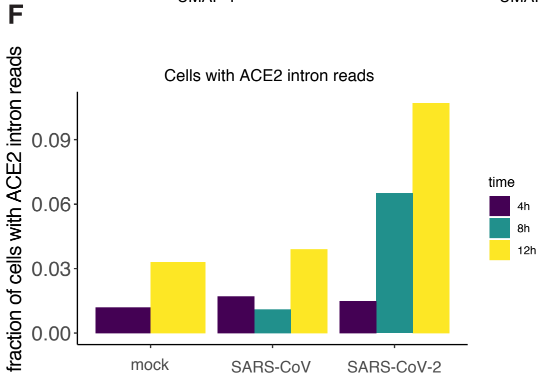

F
Cells with ACE2 intron reads
fraction of cells with ACE2 intron reads
0.09
0.06
0.03
0.00
mock
SARS-CoV
SARS-CoV-2
time
4h
8h
12h

### Figure S6

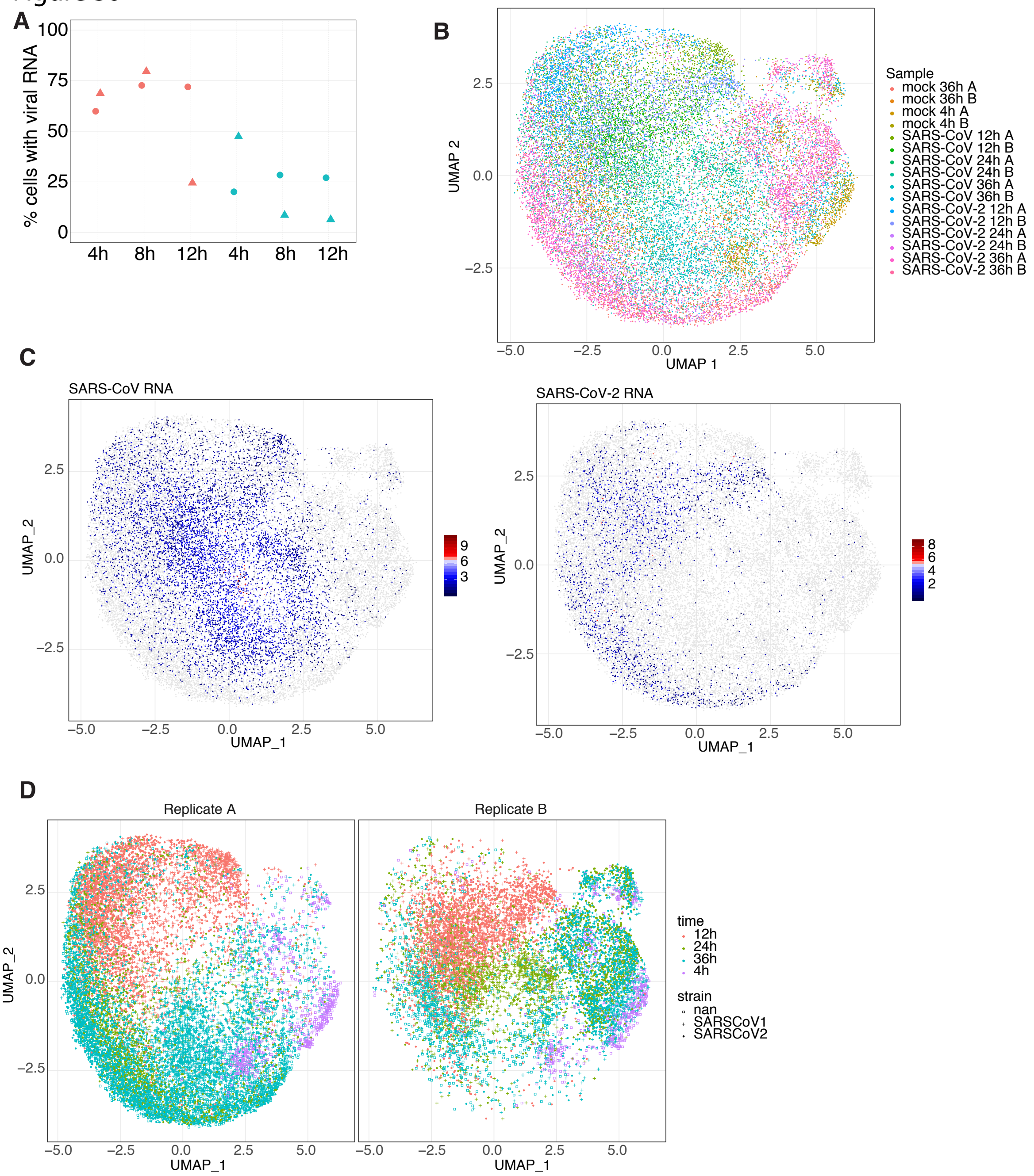
A
B
C
D
% cells with viral RNA
100
75
50
25
0
4h
8h
12h
4h
8h
12h
UMAP 2
UMAP 1
Sample
mock 36h A
mock 36h B
mock 4h A
mock 4h B
SARS-CoV 12h A
SARS-CoV 12h B
SARS-CoV 24h A
SARS-CoV 24h B
SARS-CoV 36h A
SARS-CoV 36h B
SARS-CoV-2 12h A
SARS-CoV-2 12h B
SARS-CoV-2 24h A
SARS-CoV-2 24h B
SARS-CoV-2 36h A
SARS-CoV-2 36h B
SARS-CoV RNA
SARS-CoV-2 RNA
UMAP_1
UMAP_2
Replicate A
Replicate B
time
12h
24h
36h
4h
strain
nan
SARSCoV1
SARSCoV2

Figure S7

A

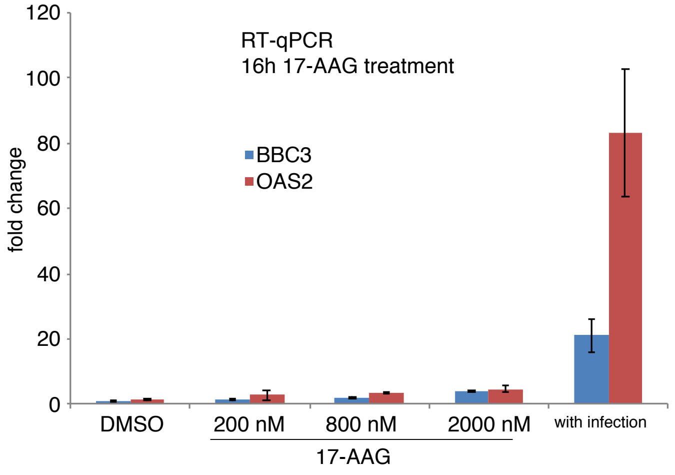
RT-qPCR
16h 17-AAG treatment
BBC3
OAS2
fold change
120
100
80
60
40
20
0
DMSO
200 nM
800 nM
2000 nM
with infection
17-AAG

B

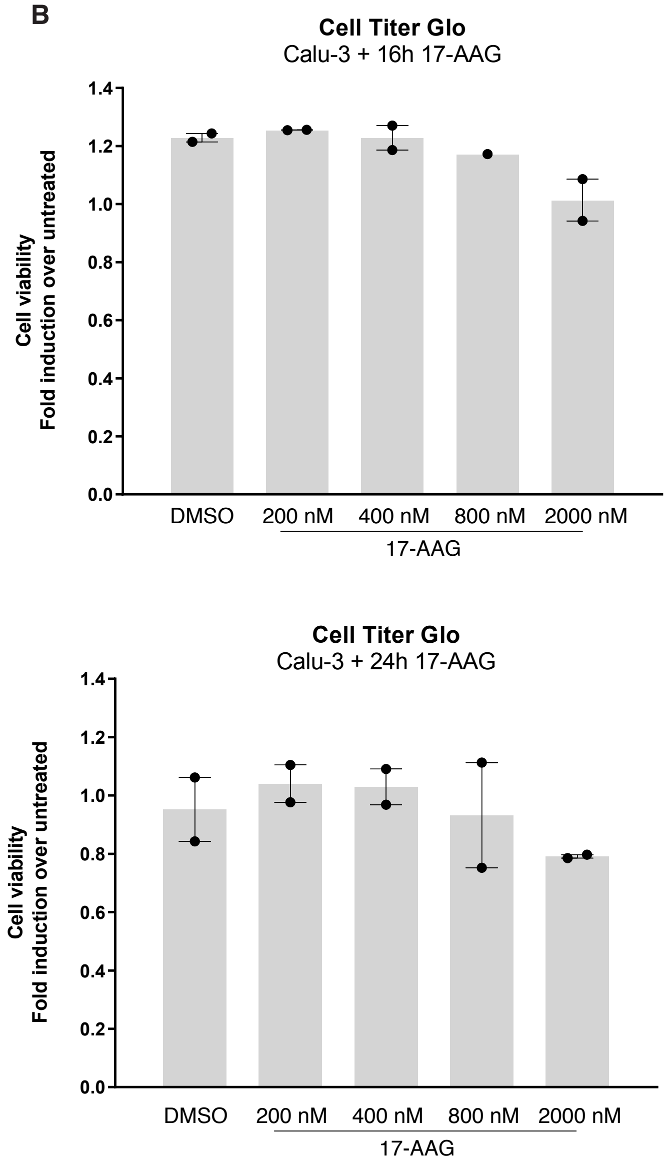
Cell Titer Glo
Calu-3 + 16h 17-AAG
Cell viability
Fold induction over untreated
1.4
1.2
1.0
0.8
0.6
0.4
0.2
0.0
DMSO
200 nM
400 nM
800 nM
2000 nM
17-AAG
Cell Titer Glo
Calu-3 + 24h 17-AAG
Cell viability
Fold induction over untreated
1.4
1.2
1.0
0.8
0.6
0.4
0.2
0.0
DMSO
200 nM
400 nM
800 nM
2000 nM
17-AAG
